# Discovery, Characterization, and Bioactivity of the Achromonodins: Lasso Peptides Encoded by *Achromobacter*

**DOI:** 10.1101/2023.06.21.545946

**Authors:** Drew V. Carson, Yi Zhang, Larry So, Wai Ling Cheung-Lee, Alexis Jaramillo Cartagena, Seth A. Darst, A. James Link

## Abstract

Through genome mining efforts, we discovered two lasso peptide biosynthetic gene clusters (BGCs) within two different species of *Achromobacter*, a genus that contains pathogenic organisms that can infect patients with cystic fibrosis. Using gene-refactored BGCs in *E. coli*, we heterologously expressed two lasso peptides, which we named achromonodin-1 and achromonodin-2. Achromonodin-1 is naturally encoded by certain isolates from the sputum of patients with cystic fibrosis. We solve the NMR structure of achromonodin-1, demonstrating that it is a threaded lasso peptide with a large loop and short tail structure, reminiscent of previously characterized lasso peptides that inhibit RNA polymerase (RNAP). We then show that achromonodin-1 inhibits RNAP *in vitro* and has potent but narrow-spectrum activity towards *Achromobacter pulmonis*, another isolate from the sputum of a cystic fibrosis patient. Our efforts expand the repertoire of antimicrobial lasso peptides and provide insights into how *Achromobacter* isolates from certain ecological niches may interact with each other.

## Introduction

Species of the genus *Achromobacter* have been causative agents of nosocomial infections and are especially prone to infect patients with cystic fibrosis.^1^ These Gram-negative pathogens, with *Achromobacter xylosoxidans* as the most clinically relevant,^2^ worryingly display intrinsic resistance to commonly prescribed antibiotic classes such as aminoglycosides,^3^ and some strains can even resist the powerful, last-resort carbapenem antibiotics.^3, 4^ To counteract the rise of antibiotic resistant pathogens, new treatments must be developed.

One compound class with demonstrated antimicrobial activity is the lasso peptides,^5^ which are characterized by their threaded [1]rotaxane structure.^6, 7^ Lasso peptide mechanisms of action range from inhibition of RNA polymerase (RNAP),^8, 9, 10^ disruption of proteolytic complexes,^11^ and inhibition of cell wall biosynthesis.^12^ We recently characterized the lasso peptide ubonodin from *Burkholderia ubonensis* and found that this lasso peptide inhibited RNAP *in vitro* and had potent antimicrobial activity against other *Burkholderia* strains.^13, 14, 15^ The discriminating factor for susceptibility was the presence of a TonB-dependent siderophore receptor, PupB.^14^ We hypothesized that other lasso peptides with structures similar to ubonodin may also serve as RNAP-inhibiting antimicrobial candidates, and we focused our efforts in finding promising candidates from the *Achromobacter* genus, a phylogenetic relative of the *Burkholderia* genus. Here we describe the heterologous expression of two lasso peptides, achromonodin-1 and achromonodin-2. We determined a solution NMR structure of achromonodin-1, demonstrated that it inhibits RNAP *in vitro*, and showed that it has potent, focused activity against an *Achromobacter* strain.

## Results and Discussion

### Genome Mining Enables Discovery of New Lasso Peptides from *Achromobacter*

Using our lasso peptide genome mining algorithm,^16^ we discovered two lasso peptide biosynthetic gene clusters (BGCs): one from a strain of *Achromobacter xylosoxidans* isolated from the sputum of a cystic fibrosis patient (found on sequenced contig from GenBank accession CYTQ01000003), and one from *Achromobacter* sp. RTa, a strain isolated from termite gut and rumen fluid (found on sequenced contig from NCBI RefSeq accession NZ_JPYO01000020). We named the putative lasso peptides from these clusters achromonodin-1 and achromonodin-2, respectively, since the peptides are native to *Achromobacter* species and the Latin root *nodum* signifies knot. We later identified BGCs responsible for producing achromonodin-1 in three other strains, including from other clinical isolates from sputum of cystic fibrosis patients^17^ and an environmental isolate from a hospital setting.

The native BGCs for both achromonodin-1 and achromonodin-2, shown in Figure 1A, contain an *A* gene, encoding the lasso peptide precursor, a *B* gene, encoding the protease responsible for producing the core peptide of the precursor, a *C* gene, encoding the cyclase that folds the core peptide and installs the key isopeptide bond, and a *D* gene, encoding the ATP-binding cassette transporter responsible for exporting the resulting lasso peptide.^18^ The genes appear in a single operon, similar to most other proteobacterial lasso peptide clusters.^19^ The presence of the dedicated lasso peptide transporter indicated that these lasso peptides may serve as antimicrobial agents to aid the host species in competing with nearby bacteria for resources, with the transporter functioning as an immunity factor.^18, 20^

**Figure 1.**
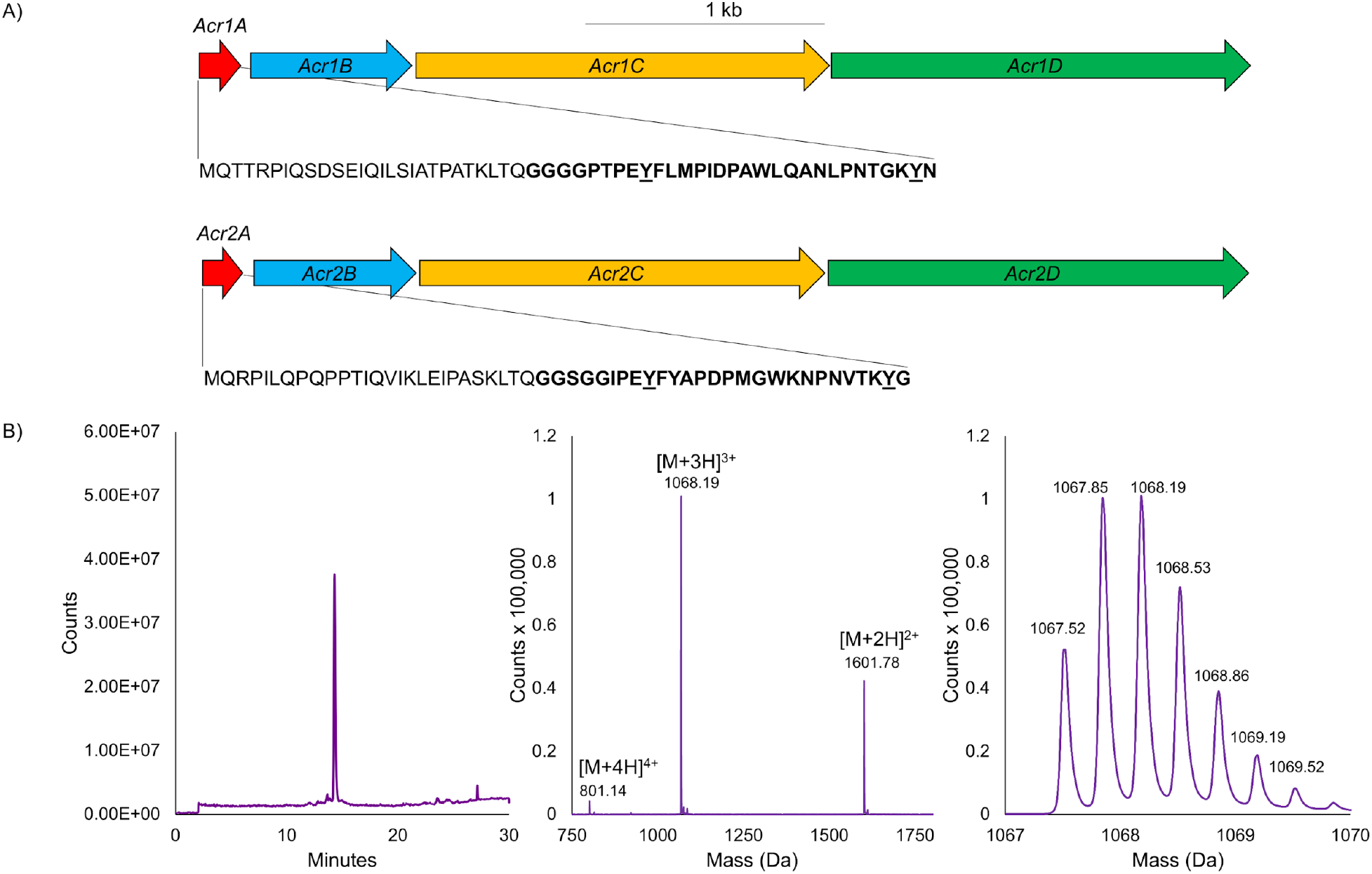
A) Achromonodin-1 and achromonodin-2 biosynthetic gene cluster organization. The amino acid sequences for the precursors are explicitly shown. The core peptide sequence is bolded, with the important tyrosine residues underlined. B) Achromonodin-1 can be heterologously expressed in *E. coli*. Left: LC-MS trace of purified achromonodin-1. Middle: Region of mass spectrum of purified achromonodin-1 where the +4, +3, and +2 charge-states appear. Right: A close-up of the mass spectrum of the +3 charge-state is displayed. Expected +3 monoisotopic m/z of achromonodin-1 is 1067.52.

In the sequences of the expected core peptides, we observed a Tyr directly after the likely eight-membered ring in both achromonodin-1 and achromonodin-2 (Tyr9) (Figure 1A). Tyr9 of the antimicrobial lasso peptide microcin J25 displayed crucial hydrogen bonding interactions with RNAP in a co-crystal structure^10^ and is also present in the potent RNAP-inhibiting lasso peptides ubonodin^13^ and citrocin.^21^ The achromonodins also include a tyrosine in the penultimate position of the core peptide, which is well-conserved across RNAP-inhibiting lasso peptides.^13, 21, 22, 23^ Due to these characteristics, we prioritized these genome mining hits for production and characterization with the hope that these would also possess RNAP-inhibiting (and antimicrobial) activity.

BlastP searches conducted on the precursor peptides from both BGCs revealed two other putative lasso peptide gene clusters from *Achromobacter* with high similarity to the achromonodins, especially achromonodin-2 (Figure S1). The strains that harbor these achromonodin-2 like BGCs are both from microbiomes; one from *C. elegans*^24^ and one from the GI tract of a goat.^25^ All of these peptides harbor the two key Tyr residues discussed above.

### Heterologous Expression of Achromonodins in *E. coli*

With many BGCs often remaining silent in native species under laboratory conditions,^26, 27^ we opted for a heterologous expression strategy in *E. coli*, which has worked well for proteobacterial lasso peptides.^28, 29, 30^ We were also unable to acquire the native producer strains for either achromonodin-1 or -2. We used codon-optimized gBlocks and oligonucleotides to assemble refactored BGCs for production of each peptide. Briefly, each precursor was placed under an isopropyl ß-D-1-thiogalactopyranoside (IPTG)-inducible T5 promoter in the pQE-80 vector. The rest of the BGC (*B, C*, and *D* genes), for both cases, was placed under the control of the constitutive promoter for the microcin J25 *BCD* operon, as has been performed previously.^13, 21, 22, 28^

We expressed these constructs with IPTG induction in *E. coli* BL-21 in M9 minimal media overnight. Due to the presence of the *D* gene, we investigated the supernatant for production of each lasso peptide since the D protein exports the lasso peptide upon production in the cell (Figure S2). While achromonodin-2 was produced and detected via LC-MS analysis of the supernatant (Figure S3), it was produced at low levels and proved challenging to isolate via HPLC purification. Thus we focused downstream efforts and subsequent characterization on achromonodin-1, as this peptide expressed at a higher level and was more readily purified to homogeneity. After two rounds of HPLC purification, achromonodin-1 was isolated at a yield of ∼0.5 mg/L culture (Figure 1B, Figure S4).

### NMR Structure of Achromonodin-1

With purified achromonodin-1 in hand, we sought to characterize its structure through 2D NMR to verify the expected threaded nature of the peptide as well as determine the steric locks (bulky amino acids that straddle the ring and prevent unthreading^31^). Due to solubility limits in water, we re-suspended the lyophilized lasso peptide in CD_3_OH for NMR analysis (concentration 10 mg/mL or ∼3.1 mM). We collected several spectra, including a TOCSY spectrum, a COSY spectrum, a ^1^H-^13^C HSQC spectrum, and two NOESY spectra at different mixing times (700 ms and 150 ms) (Figures S5-S9). We used the long mixing time of the 700 ms NOESY to aid in chemical shift assignments for each residue of the 30 aa peptide. We next integrated the NOE peaks from the NOESY spectrum (mixing time 150 ms) and used these constraints, as well as our shift assignments (Table S1) and explicit distance restraints (Table S2) around the isopeptide bond, as inputs into CYANA 2.1 to generate 20 lowest energy model structures of the peptide.

The structure of achromonodin-1 (Figure 2A) contains an 8-membered macrolactam ring between Gly1 and Glu8, with the upper and lower steric locks demonstrated to be Lys28 and Tyr29 (Table S3). Lysine is rather rare as a steric lock, as most steric locks for lasso peptides tend to be bulkier amino acids such as phenylalanine, tryptophan, tyrosine, and arginine.^32^ However, we have also observed lysine as a steric lock for the recently characterized lasso peptide lihuanodin,^33^ and lysine-17 acts as a secondary steric lock in benenodin-1, allowing for a second conformer.^34^ The NMR structure of achromonodin-1 displays an unprecedented 20 aa loop of the lasso peptide, superseding ubonodin as the lasso peptide with the longest loop experimentally verified thus far. Among experimentally verified lasso peptides, only pandonodin,^35^ at 33 aa, is larger than the 30 aa achromonodin-1 (Figure 2B).

**Figure 2.**
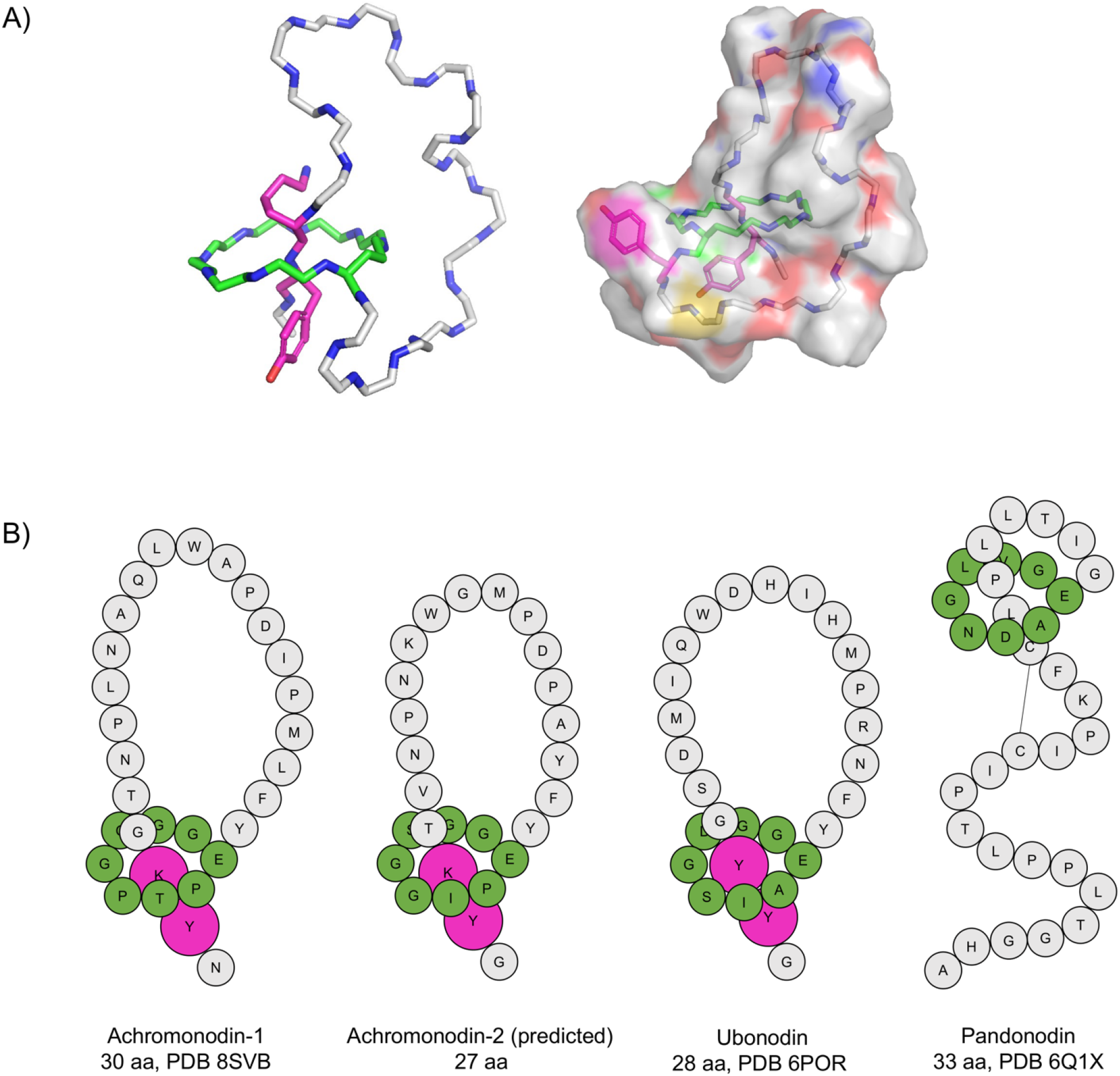
Achromonodin-1 structure and comparison to other lasso peptides. A) Left: Representative achromonodin-1 NMR structure with sidechains of steric lock residues Lys28 and Tyr29 shown in pink. Ring residue carbons are colored green, while loop and tail residue carbons are colored gray. Nitrogen atoms are colored blue and oxygen atoms are colored red. Right: Space-filling model of representative achromonodin-1 NMR structure. Color scheme is the same as the model shown on the left. Sulfur from Met12 is yellow. The sidechains of Tyr9 and Tyr29, two residues likely involved in RNAP binding, are in pink. The structure of achromonodin-1 has been deposited to the Protein Data Bank under PDB code 8SVB. B) Cartoon representation of achromonodin-1, achromonodin-2, ubonodin, and pandonodin. The structure of achromonodin-2 is predicted based on its sequence similarities to achromonodin-1 and ubonodin. For achromonodin-1, achromonodin-2, and ubonodin, the lasso peptides share similar structures, with large loops and short tails. In these representations, ring residues are shaded green and loop/tail residues are shaded gray. Steric lock residues for achromonodin-1, achromonodin-2, and ubonodin are shaded in pink and enlarged to denote their role in maintaining the threaded shape of the peptide. The cysteine residues in pandonodin are connected with a line to denote the presence of a disulfide bond in the peptide. Achromonodin-1 has the largest loop (20 aa) among experimentally verified lasso peptides to date. Pandonodin contrasts with the other lasso peptides shown here with its short loop, large tail structure.

The NMR structures demonstrate that achromonodin-1 adopts the large loop, short tail threaded lasso structure reminiscent of other RNAP-inhibiting lasso peptides, such as ubonodin (Figure 2B). The differences between the 20 model structures appear mainly in the positioning of the loop residues (Figure S10). We hypothesize that this loop remains fairly flexible in solution but will exhibit significant restructuring upon binding to its target RNAP, as was observed for microcin J25 in comparing its bound structure and solution structure.^10^ This flexible unstructured nature of the achromonodin-1 loop is analogous to what we observed for the flexibility of the large tail of pandonodin.^35^ To achieve a high enough concentration for NMR analysis, this structure was solved in methanol, but the structure is likely different in the peptide’s natural aqueous environment, especially in the flexible loop region.

While the loop of achromonodin-1 is larger than all previously characterized lasso peptides, its 8-membered ring, with Gly1 and Glu8 residues forming the isopeptide bond, is analogous to RNAP-inhibitors ubonodin, microcin J25, citrocin, klebsidin, and acinetodin. Achromonodin-1 is unique with its first four residues being glycine, perhaps providing some flexibility in the ring region. However this flexibility may be offset by the presence of two proline residues in the ring at positions 5 and 7. Additionally, this is the first example of an RNAP-inhibiting lasso peptide with a C-terminal asparagine, whereas most have a C-terminal glycine.^13, 21, 22, 23^

When we compared the top structures of achromonodin-1 to ubonodin, we noticed the presence of a kink in the loop regions nearby where a tryptophan residue of each peptide resides (Figure S10). These tryptophan residues (Trp18 for achromonodin-1 and Trp19 for ubonodin) are found in similar positions within the lasso peptide loops, and Trp18 is also conserved in achromonodin-2 (Figure S11, Figure S1). From these observations, we hypothesize that this Trp residue may be important for both peptides’ structure and function. In a recent study, we investigated the role of each residue of ubonodin for its importance in RNAP-inhibiting activity.^15^ While our data indicate that variants of Trp19 largely maintained RNAP-inhibiting activity, we speculate that this Trp residue in both peptides may play a key role in transport across the cell membrane into susceptible strains.

### Achromonodin-1 Inhibits RNA Polymerase *In Vitro*

After determining the structure of achromonodin-1, we next sought to test our hypothesis that it would inhibit RNAP. This was tested using an abortive transcription initiation assay with *E. coli* RNAP that was also used for testing ubonodin, microcin J25, and citrocin.^13, 21^ From this assay, we observed that achromonodin-1 inhibited the transcriptional capacity of *E. coli* RNAP at similar levels to ubonodin (Figure 3A, Figure S12). Thus, we conclude that achromonodin-1 inhibits RNAP, like other lasso peptides that harbor the two tyrosine residues discussed above.

**Figure 3.**
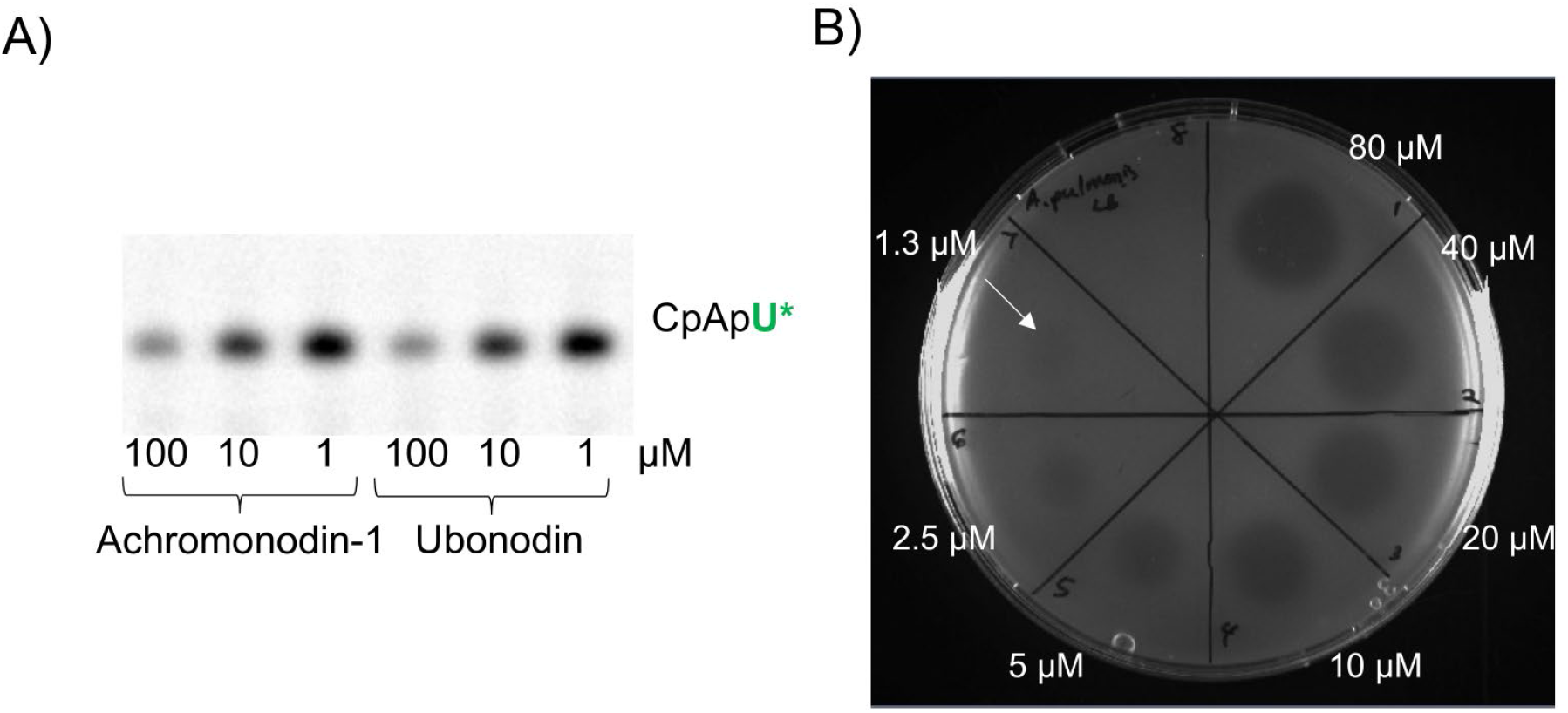
Achromonodin-1 exerts antimicrobial activity by targeting RNA polymerase (RNAP). A) *In vitro* abortive transcription initiation assay of achromonodin-1 and ubonodin against *E. coli* RNAP. Achromonodin-1 inhibits *E. coli* RNAP at similar levels to ubonodin. B) Spot-on-lawn assay showing that achromonodin-1 has antimicrobial activity against *A. pulmonis* down to 1.3 μM.

### Achromonodin-1 Displays Narrow-Spectrum Antimicrobial Activity

Knowing that achromonodin-1 can inhibit the transcriptional activity of *E. coli* RNAP, we next tested strains that achromonodin-1 may target. Since lasso peptides tend to target phylogenetically related strains, we tested a panel of three *Achromobacter* strains obtained commercially. This panel is comprised of strains that were isolated as human clinical samples: *A. pulmonis, A. insolitus*, and *A. animicus*.^36, 37^ We used a spot-on-lawn assay where LB soft agar was inoculated with the strain of interest and two-fold dilutions of the peptide were then spotted on the plate. Based on this assay, among the strains we tested we found achromonodin-1 to exclusively have activity against *A. pulmonis*, which was isolated from the sputum of a cystic fibrosis patient,^36^ at a concentration down to 1.3 μM (Figure 3B).

This narrow-spectrum antimicrobial activity is consistent with observations of other lasso peptides.^12, 13, 14, 21, 22, 23, 38^ The β and β’ of the subunits of RNA polymerase (which are the key subunits for MccJ25 binding^10^) in these species are >65% identical to their homologs in *E. coli* (Table S4), with the *Achromobacter* RNAP subunits >98% identical to each other. While we show that achromonodin-1 is active against *E. coli* RNAP, it may have higher affinity for the *Achromobacter* RNAP. We suspect that, like ubonodin, microcin J25, and microcin Y, the distinguishing factor for susceptibility is the specific transport pathway into the cell.^14, 39^

The outer membrane of Gram-negative bacteria is a major barrier for entry of many antibiotics otherwise active against Gram-positive bacteria.^40, 41, 42, 43^ While we observed no activity of achromonodin-1 against wild-type *E. coli* MG1655, we observed low micromolar (MIC of 2.5 μM) activity against the *E. coli imp4213* strain (Figure S13), a mutant strain with increased permeability of the outer membrane.^44, 45, 46^ Other RNAP-inhibiting lasso peptides cross the inner membrane via the proton gradient powered SbmA in *E. coli*^47, 48^ or the ABC transporter YddA in *Burkholderia*.^14^ *A. pulmonis* does not have any close SbmA homologs; there is a 560 aa protein annotated as an SbmA/BacA-like family transporter (accession number WP_175131624.1), but this protein is almost certainly an ABC transporter. *A. pulmonis* encodes a 614 aa YddA homolog with ∼40% identity to the *B. cepacia* protein (accession number WP_175135731.1). Achromonodin-1 may use these proteins to cross the inner membrane. Based on the results with the *E. coli imp4213* strain, we argue that the spectrum of activity of achromonodin-1 (like other RNAP-inhibiting lasso peptides) is dependent on its ability to traverse the outer membrane. Achromonodin-1 may hijack an outer membrane receptor unique to the strains that it kills, explaining the narrow-spectrum bioactivity.

## Methods

Detailed methods can be found in the supplementary material.

### Identification of Achromonodin BGCs and homologous BGCs

We used our in-house precursor centric genome mining algorithm^16^ which led us to discovery of the BGCs from *Achromobacter* species. We took interest in these because we knew that species from the *Achromobacter* genus can be a causative agent of human disease. To identify other BGCs, we conducted protein BLAST searches using the achromonodin-1 and achromonodin-2 precursors as the query.

### Cloning and Plasmid Construction

To express achromonodin-1 and achromonodin-2 in *E. coli*, we refactored the BGCs using our previously developed methods^13, 21, 28^ and designed constructs using *E. coli* codon-optimized genes. Restriction enzyme-based digestion and subsequent ligation was used to clone the BGCs into the pQE-80 vector. The final expression plasmids were named pWC132 (for achromonodin-1 expression) and pLS9 (for achromonodin-2 expression).

### Heterologous Expression and Purification of Achromonodins

To express achromonodin-1 or achromonodin-2, the corresponding expression plasmid was transformed via electroporation into *E. coli* BL21 cells. Following selection for successfully transformed colonies, the cells were cultured at large scale (500 mL culture in a 2 L flask) in M9 minimal media supplemented with amino acids. 1 mM IPTG was added to the culture when the OD600 reached about 0.2. The cultures were allowed to shake overnight at room temperature following induction.

The next day, the cultures were spun down and the supernatant was then run through a C8 column. 100% methanol was then used to elute compounds from the C8 column. We then used a rotary evaporator to remove the methanol. Dried extracts were resuspended in 50% water/50% acetonitrile. LC-MS analysis was used with the extracts to confirm expression of the lasso peptide. Two rounds of HPLC purification were used to fully isolate achromonodin-1 for further analysis.

### NMR Structure Determination

Various 2D NMR spectra were collected of purified achromonodin-1 dissolved in CD_3_OH. We then used the spectra to assign identifiable chemical shifts for hydrogen atoms and used CYANA 2.1 with the NOEs from the NOESY spectrum (150 ms mixing time) and explicit distance constraints in order to model the structure of achromonodin-1. Avogadro was then used to energy minimize each of the 20 structures of the peptide. The 20 structures were submitted to the PDB under PDB code 8SVB.

### Susceptibility Testing

To evaluate the susceptibility of the bacterial strains tested here, we used a spot-on-lawn assay. Soft agar was inoculated with a bacterial strain at 10^7^ CFU/mL, and then purified peptide was spotted at various concentrations atop the plate. *Achromobacter* strains were tested using LB soft agar and grown at 30 °C, while *E. coli* strains were tested using M63 minimal media soft agar and grown at 37 °C. Plates were incubated to allow for bacterial growth and then visualized for zones of inhibition from the spotted peptide.

## Acknowledgements

Abstract figure was created with BioRender.com. We thank István Pelczer (Princeton University NMR Facility) for assistance in NMR data acquisition of achromonodin-1. We thank Professor Mark Brynildsen (Princeton University Department of Chemical and Biological Engineering) for sharing the *E. coli imp4213* strain for susceptibility testing. This work was supported both by US National Institutes of Health (NIH) grant GM107036 to AJL as well as funding toward the Focused Research Team for Precision Antibiotics from Princeton University SEAS Funds. W.L.C-L. was supported by the Harold W. Dodds Fellowship from Princeton University. A.J.C. was supported by a Robert D. Watkins Graduate Research Fellowship from the American Society of Microbiology.

## ToC figure

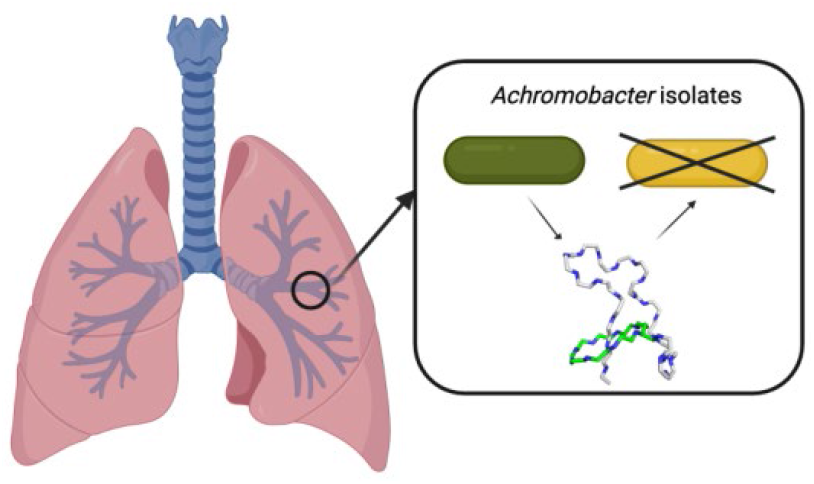

## References

1. Marion-Sanchez, K.; Pailla, K.; Olive, C.; Le Coutour, X.; Derancourt, C., Achromobacter spp. healthcare associated infections in the French West Indies: a longitudinal study from 2006 to 2016. BMC Infectious Diseases 2019, 19.

2. Menetrey, Q.; Sorlin, P.; Jumas-Bilak, E.; Chiron, R.; Dupont, C.; Marchandin, H., Achromobacter xylosoxidans and Stenotrophomonas maltophilia: Emerging Pathogens Well-Armed for Life in the Cystic Fibrosis Patients’ Lung. Genes 2021, 12.

3. Isler, B.; Kidd, T. J.; Stewart, A. G.; Harris, P.; Paterson, D. L., Achromobacter Infections and Treatment Options. Antimicrob. Agents Chemother. 2020, 64.

4. Bador, J.; Amoureux, L.; Blanc, E.; Neuwirth, C., Innate Aminoglycoside Resistance of Achromobacter xylosoxidans Is Due to AxyXY-OprZ, an RND-Type Multidrug Efflux Pump. Antimicrob. Agents Chemother. 2013, 57, 603–605.

5. Tan, S.; Moore, G.; Nodwell, J., Put a Bow on It: Knotted Antibiotics Take Center Stage. Antibiotics-Basel 2019, 8.

6. Maksimov, M. O.; Pan, S. J.; Link, A. J., Lasso peptides: structure, function, biosynthesis, and engineering. Nat. Prod. Rep. 2012, 29, 996–1006.

7. Hegemann, J. D.; Zimmermann, M.; Xie, X.; Marahiel, M. A., Lasso Peptides: An Intriguing Class of Bacterial Natural Products. Acc. Chem. Res. 2015, 48, 1909–1919.

8. Adelman, K.; Yuzenkova, J.; La Porta, A.; Zenkin, N.; Lee, J.; Lis, J. T.; Borukhov, S.; Wang, M. D.; Severinov, K., Molecular mechanism of transcription inhibition by peptide antibiotic microcin J25. Mol. Cell 2004, 14, 753–762.

9. Mukhopadhyay, J.; Sineva, E.; Knight, J.; Levy, R. M.; Ebright, R. H., Antibacterial peptide microcin J25 inhibits transcription by binding within and obstructing the RNA polymerase secondary channel. Mol. Cell 2004, 14, 739–751.

10. Braffman, N.; Piscotta, F. J.; Hauver, J.; Campbell, E. A.; Link, A. J.; Darst, S. A., Structural mechanism of transcription inhibition by lasso peptides microcin J25 and capistruin. Proc. Natl. Acad. Sci. U. S. A. 2019, 116, 1273–1278.

11. Gavrish, E.; Sit, C. S.; Cao, S. G.; Kandror, O.; Spoering, A.; Peoples, A.; Ling, L.; Fetterman, A.; Hughes, D.; Bissell, A.; Torrey, H.; Akopian, T.; Mueller, A.; Epstein, S.; Goldberg, A.; Clardy, J.; Lewis, K., Lassomycin, a Ribosomally Synthesized Cyclic Peptide, Kills Mycobacterium tuberculosis by Targeting the ATP-Dependent Protease ClpC1P1P2. Chem. Biol. 2014, 21, 509–518.

12. Tan, S.; Ludwig, K. C.; Muller, A.; Schneider, T.; Nodwell, J. R., The Lasso Peptide Siamycin-I Targets Lipid II at the Gram-Positive Cell Surface. ACS Chemical Biology 2019, 14, 966–974.

13. Cheung-Lee, W. L.; Parry, M. E.; Zong, C.; Cartagena, A. J.; Darst, S. A.; Connell, N. D.; Russo, R.; Link, A. J., Discovery of Ubonodin, an Antimicrobial Lasso Peptide Active against Members of the Burkholderia cepacia Complex. Chembiochem 2020, 21, 1335–1340.

14. Do, T.; Thokkadam, A.; Leach, R.; Link, A. J., Phenotype-Guided Comparative Genomics Identifies the Complete Transport Pathway of the Antimicrobial Lasso Peptide Ubonodin in Burkholderia. ACS Chemical Biology 2022, 17, 2332–2343.

15. Thokkadam, A.; Do, T.; Ran, X. C.; Brynildsen, M. P.; Yang, Z. J.; Link, A. J., High-Throughput Screen Reveals the Structure-Activity Relationship of the Antimicrobial Lasso Peptide Ubonodin. ACS Central Science 2023, 9, 540–550.

16. Maksimov, M. O.; Pelczer, I.; Link, A. J., Precursor-centric genome-mining approach for lasso peptide discovery. Proc. Natl. Acad. Sci. U. S. A. 2012, 109, 15223–15228.

17. Chalhoub, H.; Kampmeier, S.; Kahl, B. C.; Van Bambeke, F., Role of Efflux in Antibiotic Resistance of Achromobacter xylosoxidans and Achromobacter insuavis Isolates From Patients With Cystic Fibrosis. Frontiers in Microbiology 2022, 13.

18. Solbiati, J. O.; Ciaccio, M.; Farias, R. N.; Gonzalez-Pastor, J. E.; Moreno, F.; Salomon, R. A., Sequence analysis of the four plasmid genes required to produce the circular peptide antibiotic microcin J25. J. Bacteriol. 1999, 181, 2659–2662.

19. Maksimov, M. O.; Link, A. J., Discovery and Characterization of an Isopeptidase That Linearizes Lasso Peptides. J. Am. Chem. Soc. 2013, 135, 12038–12047.

20. Granato, E. T.; Meiller-Legrand, T. A.; Foster, K. R., The Evolution and Ecology of Bacterial Warfare. Curr. Biol. 2019, 29, R521–R537.

21. Cheung-Lee, W. L.; Parry, M. E.; Cartagena, A. J.; Darst, S. A.; Link, A. J., Discovery and structure of the antimicrobial lasso peptide citrocin. J. Biol. Chem. 2019, 294, 6822–6830.

22. Carson, D. V.; Patiño, M.; Elashal, H. E.; Cartagena, A. J.; Zhang, Y.; Whitley, M. E.; So, L.; Kayser-Browne, A. K.; Earl, A. M.; Bhattacharyya, R. P.; Link, A. J., Cloacaenodin, an Antimicrobial Lasso Peptide with Activity against Enterobacter. ACS Infectious Diseases 2023, 9, 111–121.

23. Metelev, M.; Arseniev, A.; Bushin, L. B.; Kuznedelov, K.; Artamonova, T. O.; Kondratenko, R.; Khodorkovskii, M.; Seyedsayamdost, M. R.; Severinov, K., Acinetodin and Klebsidin, RNA Polymerase Targeting Lasso Peptides Produced by Human Isolates of Acinetobacter gyllenbergii and Klebsiella pneumoniae. ACS Chemical Biology 2017, 12, 814–824.

24. Dirksen, P.; Marsh, S. A.; Braker, I.; Heitland, N.; Wagner, S.; Nakad, R.; Mader, S.; Petersen, C.; Kowallik, V.; Rosenstiel, P.; Felix, M. A.; Schulenburg, H., The native microbiome of the nematode Caenorhabditis elegans: gateway to a new host-microbiome model. BMC Biology 2016, 14.

25. Xie, F.; Jin, W.; Si, H. Z.; Yuan, Y.; Tao, Y.; Liu, J. H.; Wang, X. X.; Yang, C. J.; Li, Q. S.; Yan, X. T.; Lin, L. M.; Jiang, Q.; Zhang, L.; Guo, C. Z.; Greening, C.; Heller, R.; Guan, L. L.; Pope, P. B.; Tan, Z. L.; Zhu, W. Y.; Wang, M.; Qiu, Q.; Li, Z. P.; Mao, S. Y., An integrated gene catalog and over 10,000 metagenome-assembled genomes from the gastrointestinal microbiome of ruminants. Microbiome 2021, 9.

26. Scherlach, K.; Hertweck, C., Mining and unearthing hidden biosynthetic potential. Nature Communications 2021, 12.

27. Ren, H. Q.; Wang, B.; Zhao, H. M., Breaking the silence: new strategies for discovering novel natural products. Curr. Opin. Biotechnol. 2017, 48, 21–27.

28. Pan, S. J.; Cheung, W. L.; Link, A. J., Engineered gene clusters for the production of the antimicrobial peptide microcin J25. Protein Expr. Purif. 2010, 71, 200–206.

29. Kodani, S.; Unno, K., How to harness biosynthetic gene clusters of lasso peptides. J. Ind. Microbiol. Biotechnol. 2020, 47, 703–714.

30. Kunakom, S.; Eustaquio, A. S., Heterologous Production of Lasso Peptide Capistruin in a Burkholderia Host. Acs Synthetic Biology 2020, 9, 241–248.

31. Allen, C. D.; Chen, M. Y.; Trick, A. Y.; Le, D. T.; Ferguson, A. L.; Link, A. J., Thermal Unthreading of the Lasso Peptides Astexin-2 and Astexin-3. ACS Chemical Biology 2016, 11, 3043–3051.

32. Hegemann, J. D., Factors Governing the Thermal Stability of Lasso Peptides. Chembiochem 2020, 21, 7–18.

33. Cao, L.; Beiser, M.; Koos, J. D.; Orlova, M.; Elashal, H. E.; Schroder, H. V.; Link, A. J., Cellulonodin-2 and Lihuanodin: Lasso Peptides with an Aspartimide Post-Translational Modification. J. Am. Chem. Soc. 2021, 143, 11690–11702.

34. Zong, C.; Wu, M. J.; Qin, J. Z.; Link, A. J., Lasso Peptide Benenodin-1 is a Thermally Actuated [1]Rotaxane Switch. J. Am. Chem. Soc. 2017, 139, 10403–10409.

35. Cheung-Lee, W. L.; Cao, L.; Link, A. J., Pandonodin: A Proteobacterial Lasso Peptide with an Exceptionally Long C-Terminal Tail. ACS Chemical Biology 2019, 14, 2783–2792.

36. Vandamme, P.; Moore, E. R. B.; Cnockaert, M.; De Brandt, E.; Svensson-Stadler, L.; Houf, K.; Spilker, T.; LiPuma, J. J., Achromobacter animicus sp nov., Achromobacter mucicolens sp nov., Achromobacter pulmonis sp nov and Achromobacter spiritinus sp nov., from human clinical samples. Syst. Appl. Microbiol. 2013, 36, 1–10.

37. Coenye, T.; Vancanneyt, M.; Falsen, E.; Swings, J.; Vandamme, P., Achromobacter insolitus sp nov and Achromobacter spanius sp nov., from human clinical samples. International Journal of Systematic and Evolutionary Microbiology 2003, 53, 1819–1824.

38. Knappe, T. A.; Linne, U.; Zirah, S.; Rebuffat, S.; Xie, X. L.; Marahiel, M. A., Isolation and structural characterization of capistruin, a lasso peptide predicted from the genome sequence of Burkholderia thailandensis E264. J. Am. Chem. Soc. 2008, 130, 11446–11454.

39. Li, Y.; Han, Y.; Zeng, Z. W.; Li, W. J.; Feng, S. X.; Cao, W. S., Discovery and Bioactivity of the Novel Lasso Peptide Microcin Y. J. Agric. Food Chem. 2021, 69, 8758–8767.

40. Lehman, K. M.; Grabowicz, M., Countering Gram-Negative Antibiotic Resistance: Recent Progress in Disrupting the Outer Membrane with Novel Therapeutics. Antibiotics-Basel 2019, 8.

41. Cama, J.; Henney, A. M.; Winterhalter, M., Breaching the Barrier: Quantifying Antibiotic Permeability across Gram-negative Bacterial Membranes. J. Mol. Biol. 2019, 431, 3531–3546.

42. Lee, C. R.; Cho, I. H.; Jeong, B. C.; Lee, S. H., Strategies to Minimize Antibiotic Resistance. International Journal of Environmental Research and Public Health 2013, 10, 4274–4305.

43. Choi, U.; Lee, C. R., Distinct Roles of Outer Membrane Porins in Antibiotic Resistance and Membrane Integrity in Escherichia coli. Frontiers in Microbiology 2019, 10.

44. Sampson, B. A.; Misra, R.; Benson, S. A., Identification and Characterization of a New Gene of Escherichia coli K-12 Involved in Outer Membrane Permeability. Genetics 1989, 122, 491–501.

45. Wu, T.; Malinverni, J.; Ruiz, N.; Kim, S.; Silhavy, T. J.; Kahne, D., Identification of a multicomponent complex required for outer membrane biogenesis in Escherichia coli. Cell 2005, 121, 235–245.

46. Chou, W. K.; Vaikunthan, M.; Schroder, H. V.; Link, A. J.; Kim, H.; Brynildsen, M. P., Synergy Screening Identifies a Compound That Selectively Enhances the Antibacterial Activity of Nitric Oxide. Frontiers in Bioengineering and Biotechnology 2020, 8.

47. Salomon, R. A.; Farias, R. N., The Peptide Antibiotic Microcin 25 Is Imported through the TonB Pathway and the SbmA Protein. J. Bacteriol. 1995, 177, 3323–3325.

48. Ghilarov, D.; Inaba-Inoue, S.; Stepien, P.; Qu, F.; Michalczyk, E.; Pakosz, Z.; Nomura, N.; Ogasawara, S.; Walker, G. C.; Rebuffat, S.; Iwata, S.; Heddle, J. G.; Beis, K., Molecular mechanism of SbmA, a promiscuous transporter exploited by antimicrobial peptides. Science Advances 2021, 7.

