## Supporting Info for "Discovery, Characterization, and Bioactivity of the Achromonodins: Lasso Peptides Encoded by *Achromobacter*"

<u>Contents</u>

Detailed methods: Pages S2-S5

13 supplementary figures: Pages S6-S16

6 supplementary tables: Pages S17-S24

References for supplementary information: Page S25

### <u>Methods</u>

<u>Materials</u>

*E. coli* XL1-blue cells were used to clone plasmids used for the expression of achromonodins, and *E. coli* BL21 cells were used for expression of the lasso peptides. gBlocks and oligonucleotides were purchased from IDT. Codon optimization for the gBlocks was accomplished using DNAWorks.[1] For cloning, genes were PCR-amplified using Q5 DNA polymerase from New England Biolabs (NEB) and digested with restriction enzymes also purchased from NEB for cloning into the pQE-80L vector from QIAGEN. QIAGEN mini-prep kits were used to purify cloned plasmids and were sequenced verified using Genewiz/Azenta before use for expression. We acquired the following *Achromobacter* strains from DSMZ: *Achromobacter pulmonis* strain LMG 26696, *Achromobacter insolitus* strain DSM 23807 and *Achromobacter animicus* strain LMG 26690.

#### Identification of Achromonodin Biosynthetic Gene Clusters

Our in-house precursor-centric genome mining algorithm[2] was used to search for precursors with a Tyr residue after the ring and a penultimate Tyr residue. The algorithm searches for open reading frames with sequence X10-43TXXX5-7[D/E]X5-25, where X signifies any of the 20 amino acids. From our previous experience with RNA polymerase-inhibiting lasso peptides, we were specifically interested in lasso peptides with a Y directly after the acidic residue (D/E), and a Y in the penultimate position.

The achromonodin-1 precursor peptide was identified with protein accession number CUI65249.1, the sequence of which is identical to protein accession WP_165596049.1. The achromonodin-1 BGC is present in four *Achromobacter xylosoxidans* strains: strain 2016Y70914664II on contig with NCBI RefSeq NZ_JAJLQM010000005.1, strain 2017Y71057280IX on a contig with NCBI RefSeq NZ_JAJLQR010000044.1, strain 2789STDY5663426 on a contig with GenBank accession CYTQ01000003.1, and strain GD04008on a contig with NCBI RefSeq NZ_JAOEEL010000023.1. Achromonodin-2 was identified with protein accession number WP_156124238.1 on a contig with NCBI RefSeq accession NZ_JPYO01000020. Genes encoding for B, C, and D proteins were all found downstream of the precursors.

After identification of the gene clusters, the precursor sequences for achromonodin-1 and achromonodin-2 were used as query sequences for a BlastP search. Default parameters were used, and the sequences were queried against the non-redundant protein sequences database. A total of four different lasso peptide precursors were found from *Achromobacter* species (Figure S1). Genes encoding for B, C, and D proteins nearby were verified.

The sequences of the lasso peptide precursors from *Achromobacter* species were aligned using the multiple sequence alignment tool from the Clustal Omega web server.[3]

#### Cloning and Plasmid Construction

For heterologous expression of both achromonodin-1 and achromonodin-2, we first used DNAWorks[1] to codon optimize the BGCs for *E. coli* production. For each plasmid for achromonodin-1 and achromonodin-2 production, the codon-optimized *A* gene was constructed via overlap PCR from oligonucleotides. The resulting PCR product was digested and ligated into the *Eco*RI and *Hin*dIII restriction sites of pQE-80 (Table S6). The strong ribosome binding site encoded downstream of the *Eco*RI site in pQE-80 was maintained by including it in the oligonucleotide construction for the overlap PCR. For each construct, the *B, C,* and *D* genes were synthesized as gBlocks with the upstream constitutive promoter for *mcjBCD* from the native BGC of microcin J25 (Table S5). After overlap PCR amplification of the gBlocks, they were cloned into the *Nhe*I and *Nco*I site of the vector previously cloned with the precursor in the pQE-80 vector. The final plasmids used for expression were pWC132 for achromonodin-1 production and pLS9 for achromonodin-2 production, consisting of a $p_{T5}$-*A* $p_{mcjBCD}$*BCD* architecture in both cases with their respective *ABCD* genes.

**Heterologous Expression and Purification of Achromonodins**

The pWC132 (for achromonodin-1 production) and pLS9 (for achromonodin-2 production) plasmids were separately used to transform via electroporation *E. coli* BL-21 cells and plated on LB agar containing 100 µg/mL ampicillin. The agar plate was incubated at 37 °C overnight to allow for visible colony formation, from which an individual colony was then used to inoculate a 5 mL solution of LB media containing 100 µg/mL ampicillin. This culture was grown with shaking at 250 rpm at 37 °C overnight, and then used to inoculate a 500 mL culture of M9 media (supplemented with 100 µg/mL ampicillin) at a starting OD600 of 0.02. This media is composed of the following: M9 salts, 0.2% w/v glucose, 1 mM $MgSO_4$, 0.00005% w/v thiamine, each of the 20 amino acids at 0.04 g/L. The 500 mL culture was grown in a heat-controlled shaker (37 °C, 250 rpm) and when the $OD_{600}$ measured 0.2, 1 mM isopropyl-ß-D-thiogalactopyranoside (IPTG) was added to induce expression of the precursor (and thus expression of achromonodin-1 or 2). The induced cultures were grown overnight (room temperature, 250 rpm).

Following overnight expression, the cultures were spun down at 4000 x *g* for 15-20 minutes and the supernatants were retained. A 6 mL Strata C8 column was used for further purification of the peptides in the supernatant. 6 mL of methanol was first flowed through the column followed by 12 mL of DI water. The supernatant was then allowed to flow through the column. 12 mL of DI water was then flowed through before elution with 6 mL of methanol. This methanol elution was retained and rotovapped. The dried eluent was then re-suspended in a 1:1 solution of water:acetonitrile (500 µL per 500 mL culture).

**LC-MS Analysis**

The supernatant extract was analyzed with LC-MS to detect the presence of achromonodin-1 or achromonodin-2. The LC-MS used was an Agilent 6530 QTOF that was connected to an Agilent 1260 LC. The instrument was used with electrospray ionization in positive ion mode. The column used was an Agilent ZORBAX StableBond 300 C18, 2.1 x 50 mm, 3.5 µm. The data was analyzed with Agilent MassHunter Qualitative Analysis software. Typically, 2-3 µL of the supernatant extract was injected into the column for measurement. The LC-MS was run at a flowrate of 0.5 mL/min. The solvents used were water and acetonitrile, both supplemented with 0.1% formic acid. The gradient used was as following: 0-1 minute of 90% water/10% acetonitrile, 1-20 minutes a linear gradient from 90% water/10% acetonitrile to 50% water/50% acetonitrile, 20-25 minutes a linear gradient from 50% water/50% acetonitrile to 10% water/90% acetonitrile.

**HPLC Purification**

Achromonodin-1 was pursued for further purification via RP-HPLC. The instrument used for this was an Agilent 1200 series HPLC. The column used was an Agilent ZORBAX StableBond 300 C18, 9.4 x 250 mm, 5 µm. The HPLC was run at a flowrate of 4 mL/min using the solvents water and acetonitrile, both supplemented with 0.1% trifluoroacetic acid.

For the first round of HPLC purification, the following gradient was used: 0-1 minute of 90% water/10% acetonitrile, 1-20 minutes a linear gradient from 90% water/10% acetonitrile to 50% water/50% acetonitrile, 25-29 minutes a linear gradient from 50% water/50% acetonitrile to 10% water/90% acetonitrile. The peak at 18.5 minutes was collected, confirmed to contain achromonodin-1 via LC-MS analysis, frozen at -80 °C, and then lyophilized. The lyophilized powder was then resuspended in 1:1 water:acetonitrile for further purification.

For the second round of HPLC purification, the following gradient was used: 0-1 minute of 90% water/10% acetonitrile, 1-3 minutes a linear gradient from 90% water/10% acetonitrile to 70% water/30% acetonitrile, 3-25 minutes a linear gradient from 70% water/30% acetonitrile to 60% water/40% acetonitrile, 25-28 minutes a linear gradient from 70% water/30% acetonitrile to 10% water/90% acetonitrile. The peak eluting at ~18.7 minutes was collected, frozen at -80 °C, and lyophilized. The resulting powder was used for further analysis.

**NMR Data Collection**

NMR data was collected at the NMR Facilities at the Princeton University Department of Chemistry on a Bruker Avance III HD 800-MHz NMR spectrometer. Lyophilized achromonodin-1 was dissolved at a concentration of 10 mg/mL in $CD_3OH$. NMR data was collected at a temperature of 22 °C (295 K). The following NMR spectra were collected: $^1$H-$^1$H TOCSY at 80 ms mixing time, $^1$H-$^1$H NOESY at 700 ms mixing time, $^1$H-$^1$H NOESY at 150 ms mixing time, $^1$H-$^1$H gCOSY, and $^1$H-$^{13}$C HSQC.

**NMR Analysis**

MestreNova software (MestreLab) was used for data processing and analysis of NMR data. The TOCSY, NOESY with 700 ms mixing time, gCOSY, and HSQC were used together to assist in chemical shift assignments of the residues of achromonodin-1. NOEs were manually picked and integrated from the NOESY (150 ms mixing time). The chemical shift assignments, NOE peaks and integrations, and explicit distance restraints around the isopeptide bond between Gly1 and Glu8 were used as inputs for CYANA 2.1.

CYANA 2.1[4] was used for seven cycles of automated NOE peak assignments and structure calculations. 100 initial structures were calculated, and after a final structural calculation, the 20 lowest energy structures were chosen as models. These structures were energy minimized using Avogadro.[5] In Avogadro, we used the force field MMFF94 and the steepest descent algorithm. The structures were submitted to the Protein Data Bank under PDB code 8SVB and to the Biological Magnetic Resonance Data Bank under BMRB code 31086.

### RNA Polymerase Inhibition Assay

Achromonodin-1 was tested alongside with the RNAP-inhibiting lasso peptides microcin J25, citrocin, and ubonodin in an assay that measures abortive transcription; the conditions of this assay are described in our reports of citrocin[6] and ubonodin.[7] Briefly, various concentrations of the peptides were tested in their ability to inhibit transcription by *E. coli* RNAP which could be detected by incorporation of radioactive uracil onto a growing RNA chain.

### Spot-on-Lawn Assay

A spot-on-lawn assay that has previously been used in our lab was used to test antimicrobial activity of achromonodin-1. *Achromobacter* strains to be tested were streaked out on LB agar and incubated at 30 °C for two nights until visible colony formation was observed that was large enough to be picked. A colony was then used to inoculate 5 mL of LB which was then grown overnight at 30 °C with shaking at 250 rpm. The next morning, the overnight culture was diluted 1:100 to inoculate 5 mL of LB, which was then grown at 30 °C with shaking at 250 rpm. Once the cells reached an OD of about 0.4-0.6, $10^8$ total CFU (assuming OD of 1 is $10^9$ CFU/mL) was added to 10 mL of melted LB soft agar (6.5% agar). The inoculated agar was poured onto a plate containing a base of 10 mL of solid M63 hard agar (components: 1X M63 salts, 0.2% glucose, 1 mM $MgSO_4$, and 0.0005% w/v thiamine). Once the soft agar dried, 10 µL of achromonodin-1 (in water) at varying concentrations were spotted on top of the agar and allowed to dry. The agar plates were then incubated at 30 °C for two nights and imaged to view spots of inhibition. The strain was deemed susceptible to achromonodin-1 if visible spots of inhibition could be observed, and the MIC is defined as the last dilution where a spot was visible.

For the *E. coli* strains that were tested here using this assay, the bacteria were grown at 37 °C instead of 30 °C. The agar used for these *E. coli* strains was 10 mL of soft M63 agar (components: 1X M63 salts, 0.2% glucose, 1 mM $MgSO_4$, 0.0005% thiamine, 40 mg/L of each of the 20 common amino acids, 6.5% agar). The agar plates for the spot-on-lawn assay for the *E. coli* strains were only incubated for one night as opposed to two nights for the *Achromobacter* species.

### Data Availability

The structure of achromonodin-1 has been deposited to the PDB under under PDB code 8SVB and to the Biological Magnetic Resonance Data Bank under BMRB code 31086. Raw data used to solve the NMR structure have been uploaded to the DataSpace at Princeton University (doi.org/10.34770/mzkj-qh70).

```
                                                   -25  -20  -15  -10  -5    1   5   10   15   20   25   30
                                                  ..|....|....|....|....|....|...|....|....|....|....|....|
              Acr1A WP_165596049.1 and CUI65249.1 MQTTRPIQSDSEIQILSIATPATKLTQGGGGPTPEYFLMPIDPAWLQANLPNTGKYN
                             Acr2A WP_156124238.1 MQRPILQPQPPTIQVIKLEIPASKLTQGGSGGIPEYFYAPDPMGWKNPNVTKYG---
Achromobacter sp. isolate RGIG2230 MBQ2647785.1   MQQRTSKPQEQPIRVIKLDIPASKLTQGGSGGIPEYFYAPDPMNWNNPNVTKYG---
Achromobacter spanius strain MYb73 WP_158685952.1 MQQHVIKSQVQAIRVIKLAVPASQLTQGGSGGIPEYFYAPDPMNFRNPNVTKYG
                                                  **      .   *:::.:  **::*****.*  ****  *    : : *: : *
```

**Figure S1.** Alignment of putative lasso peptide precursors found in *Achromobacter* species. The putative leader peptide is shown in red text. Achromonodins-1 (Acr1A) and -2 (Acr2A) are the first two entries. The isopeptide bond in all four of the putative lasso peptides is formed between the N-terminal glycine and a glutamic acid at position 8.

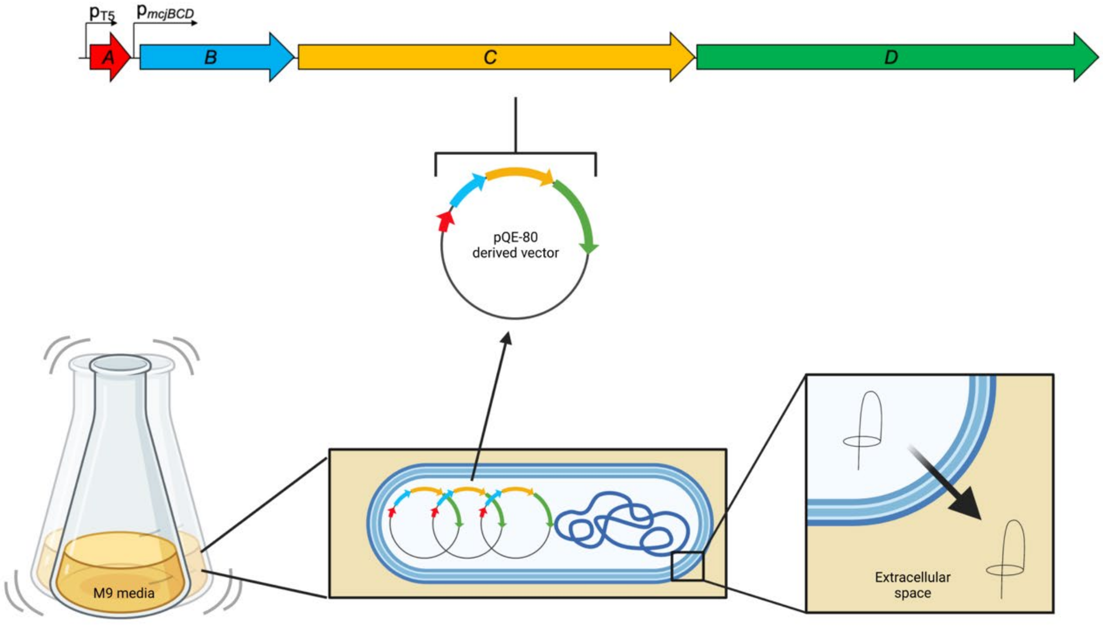


**Figure S2.** Heterologous expression strategy for producing the achromonodin lasso peptides. The gene cluster was refactored as shown at the top. The *A* gene was placed under an IPTG-inducible T5 promoter, and the rest of the gene cluster was placed under the constitutive promoter for the microcin J25 *BCD* genes. These were cloned into the pQE-80 plasmid backbone. The cells were expressed on a large scale in *E. coli* BL-21 cells in M9 media. The lasso peptides were exported to the supernatant due to the presence of the *D* gene. Created with BioRender.com.

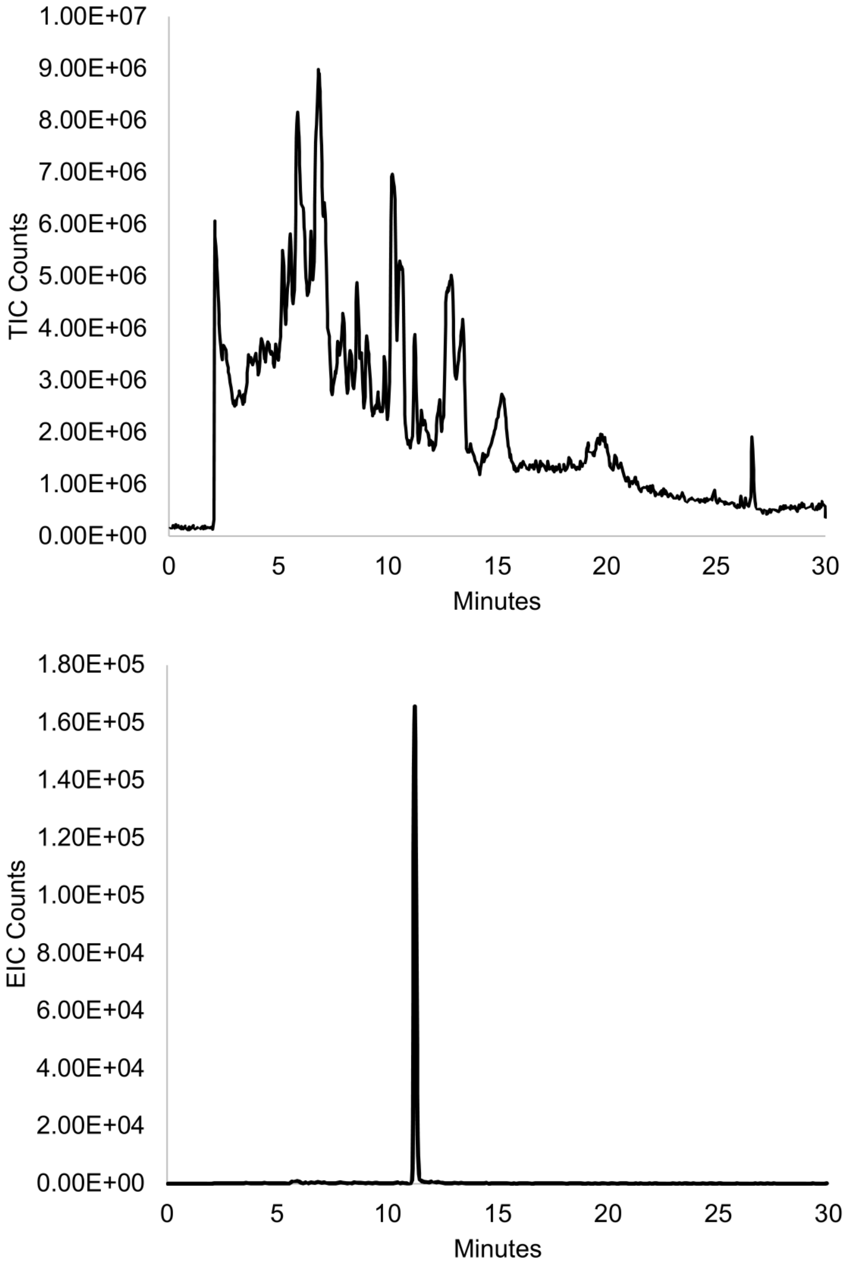


**Figure S3.** Achromonodin-2 can be successfully expressed in *E. coli.* Top: Total ion current (TIC) chromatogram of supernatant extract for *E. coli* expressing achromonodin-2. Bottom: Extracted ion current (EIC) chromatogram of supernatant extract. We extracted for the expected +3 and +2 monoisotopic charge-state of the achromonodin-2 mass, within a 50-ppm margin of error. The expected monoisotopic +3 charge-state is 962.1182 Da and the expected +2 monoisotopic charge-state is 1442.6736 Da.

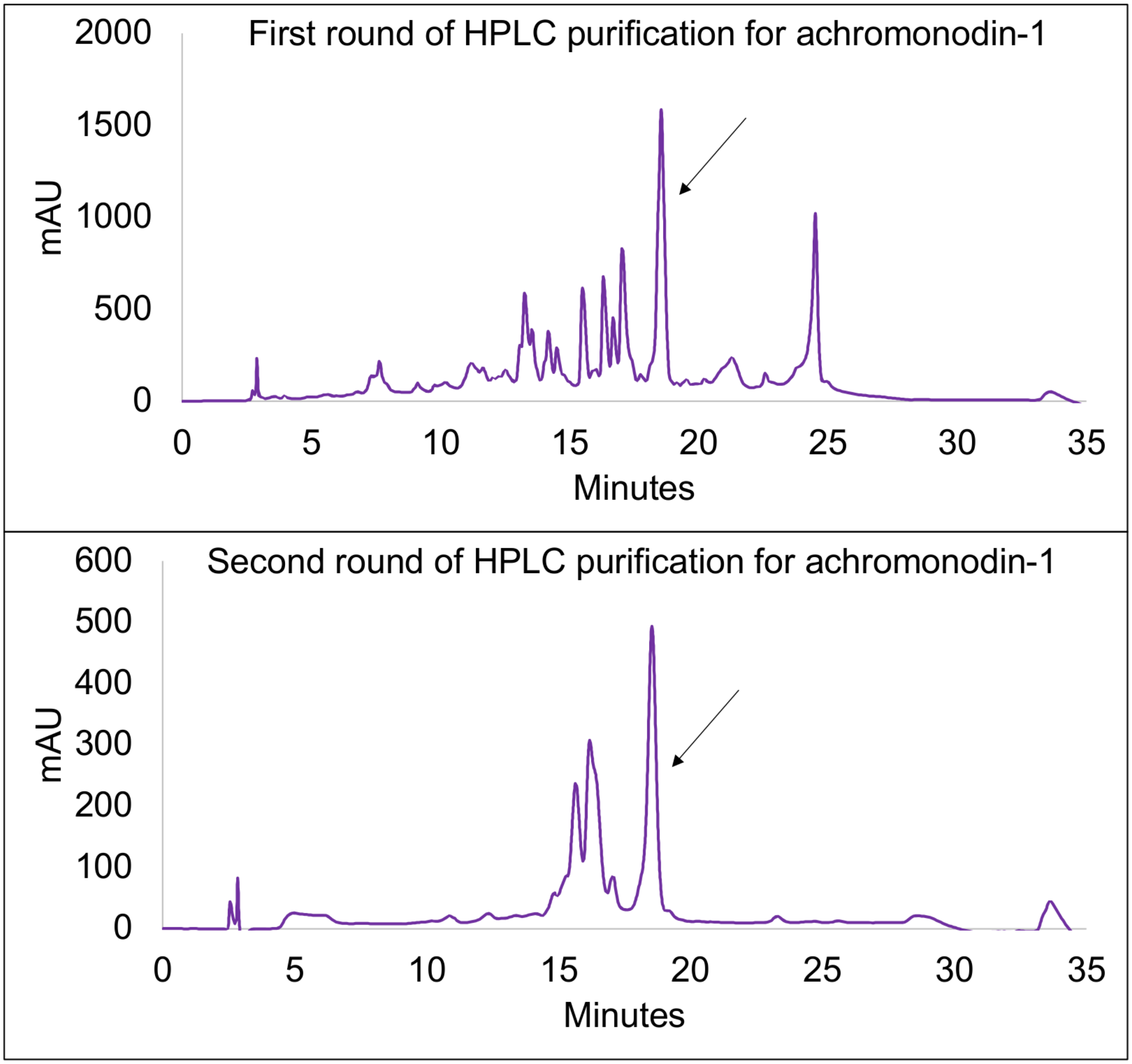


**Figure S4.** HPLC traces for achromonodin-1 purification. Arrows point to the peak that was collected from the run. Top: first round of HPLC purification using supernatant extract from *E. coli* that had expressed achromonodin-1. The conditions for this first round of HPLC purification are described in the Methods section of the SI. Bottom: second round of HPLC purification using the collected peak from the first round. The conditions for this second round of HPLC purification are also described in the Methods section of the SI.

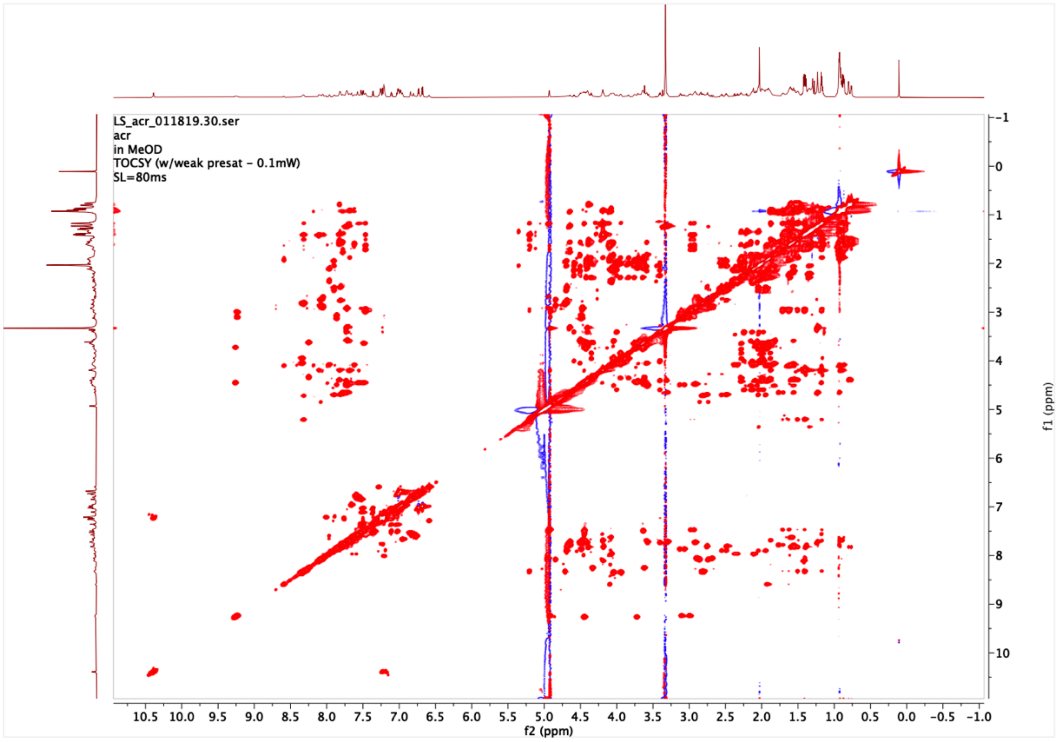


**Figure S5.** TOCSY spectrum obtained for achromonodin-1 in $CD_3OH$.*

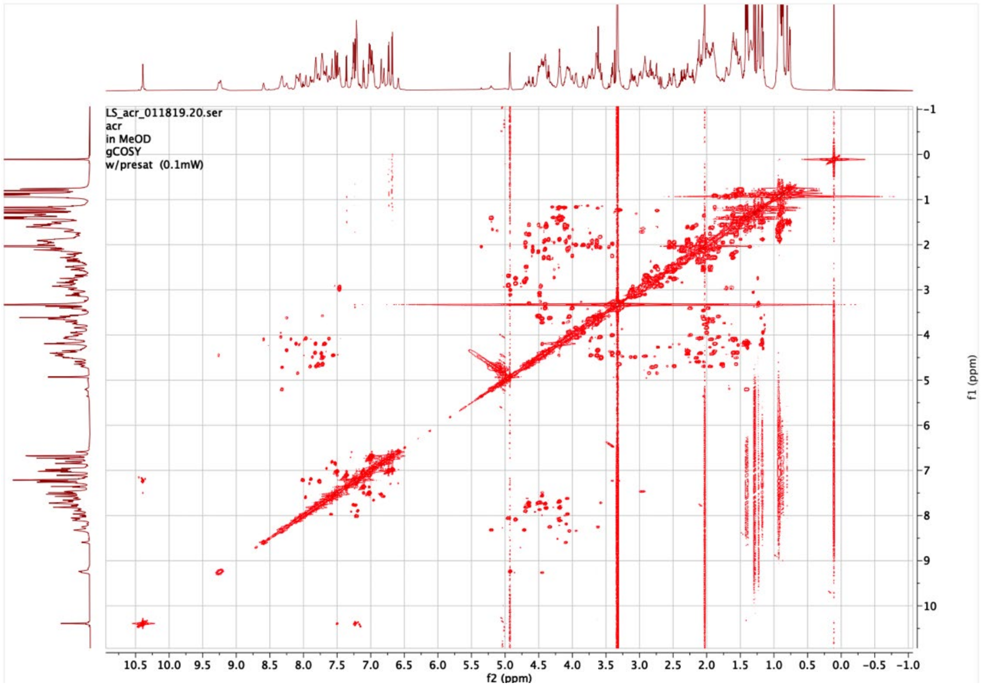


**Figure S6.** gCOSY spectrum obtained for achromonodin-1 in $CD_3OH$.*

*Note that the spectrum has MeOD in the top left corner, but this is an error; it should say $CD_3OH$.

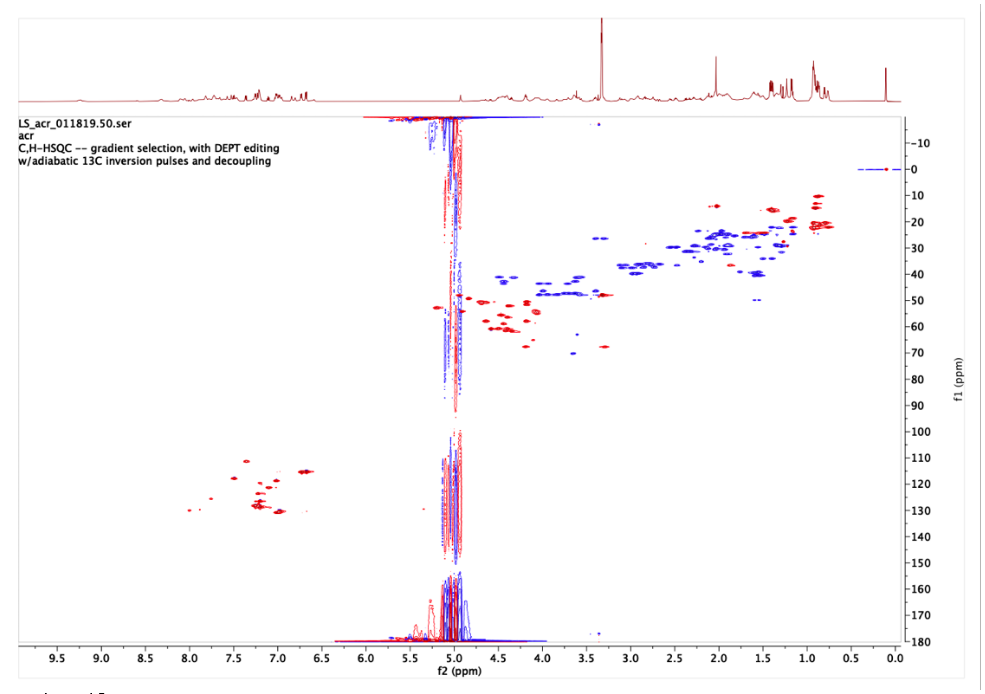


**Figure S7.** $^1H$-$^{13}C$ HSQC spectrum obtained for achromonodin-1 in $CD_3OH$. Details of the conditions for the experiment are shown in the top left corner of the spectrum.

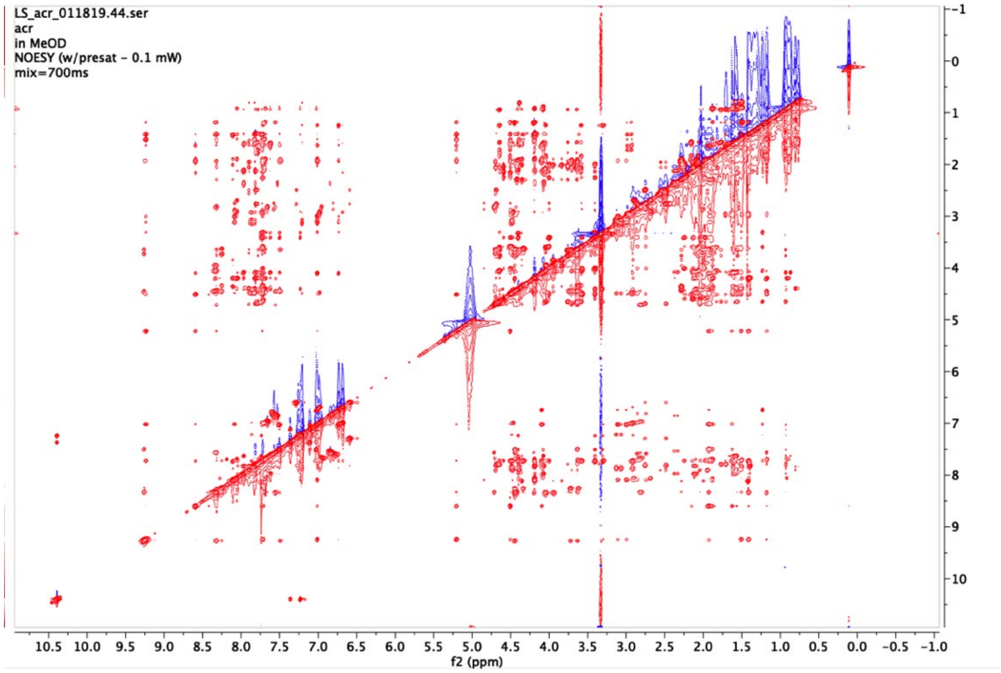


**Figure S8.** NOESY spectrum obtained for achromonodin-1 in $CD_3OH$.* Mixing time of 700 ms.

*Note that the spectrum has MeOD in the top left corner, but this is an error; it should say $CD_3OH$.

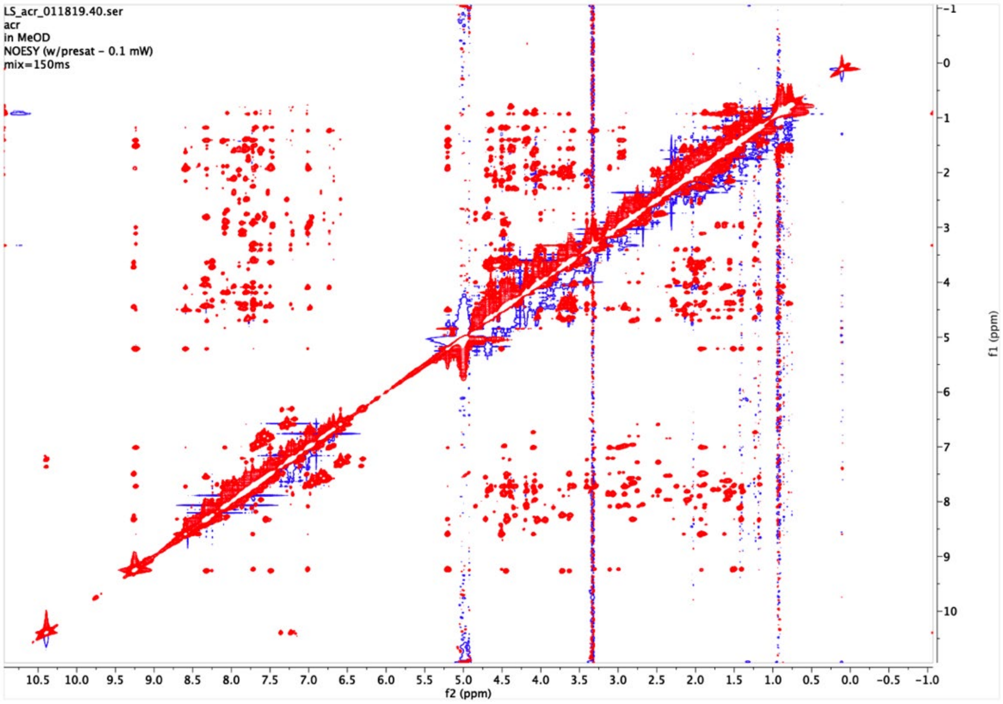


**Figure S9.** NOESY spectrum obtained for achromonodin-1 in $CD_3OH$.* Mixing time of 150 ms.*

*Note that the spectrum has MeOD in the top left corner, but this is an error. It should say $CD_3OH$.

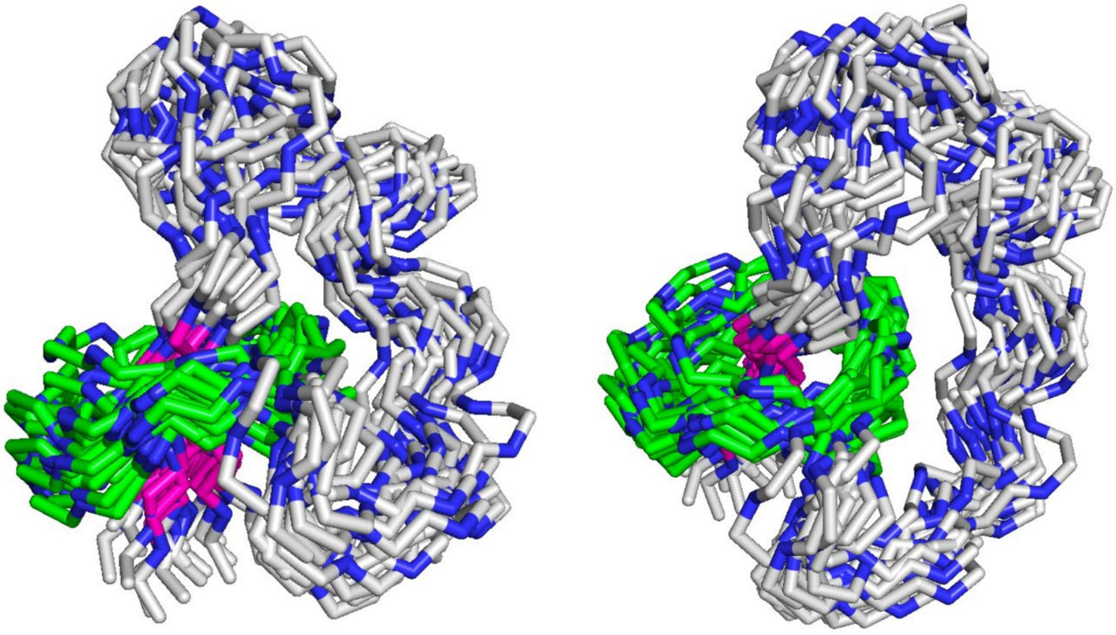

**Figure S10.** Overlay of 20 lowest-energy structures of achromonodin-1 as modeled by CYANA 2.1. Two different views are shown. We expect that the large loop is likely flexible in solution, and that the peptide adopts a different structure in water rather than this structure solved in methanol. These structures have been deposited to the Protein Data Bank under PDB code 8SVB.

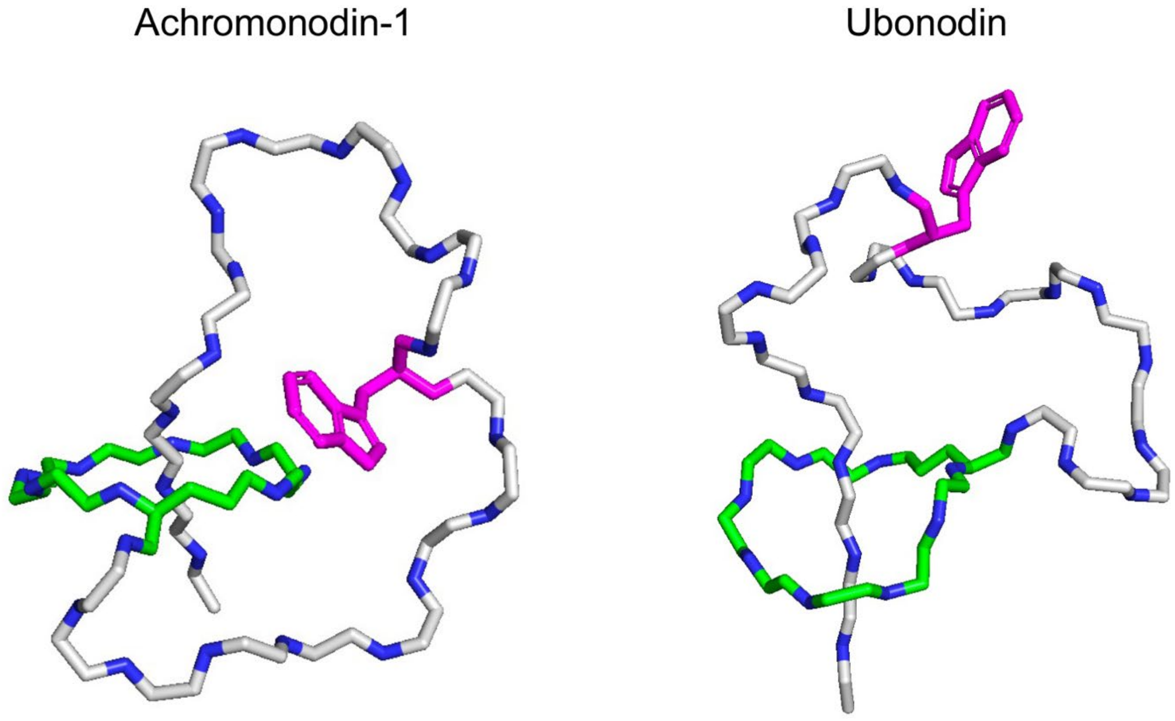


```
Ubonodin       GGDGSIAEYFNRPMHIHDW---QIMDSGYYG
               ||.|...|||..|:. ..|   .:.::|.|.
Achromonodin-1 GGGGPTPEYFLMPID-PAWLQANLPNTGKYN
```

**Figure S11.** Both achromonodin-1 and ubonodin have a tryptophan in their loop regions. This tryptophan may play a role in the structure or function of the peptides, particularly to transport into the cell. Top: The top structures of both achromonodin-1 and ubonodin are shown. Ubonodin drawn from PDB 6POR. Ring residue carbons are colored green, while loop and tail residue carbons are colored gray. Nitrogen atoms are colored blue. Trp18 of achromonodin-1 and Trp19 of ubonodin are colored in pink, and the sidechain is displayed. Bottom: Sequence alignment (completed using the EMBOSS Needle pairwise sequence alignment tool on the web server[3]) of achromonodin-1 and ubonodin. The conserved Trp residue is bolded.

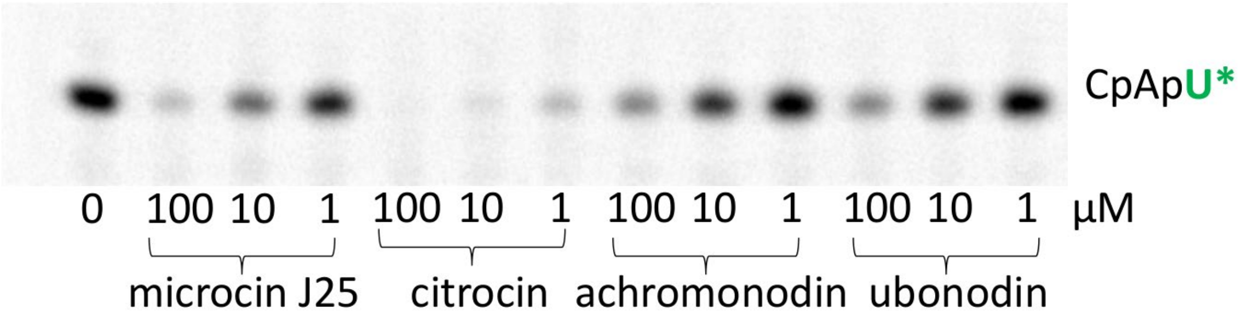


**Figure S12.** Gel for abortive transcription initiation assay showing the four different peptides tested using *E. coli* RNAP. Achromonodin in the figure refers to achromonodin-1. Achromonodin-1 behaves similarly to ubonodin in this assay. Portions of this figure were shown in previous publications for citrocin[6] and ubonodin[7].

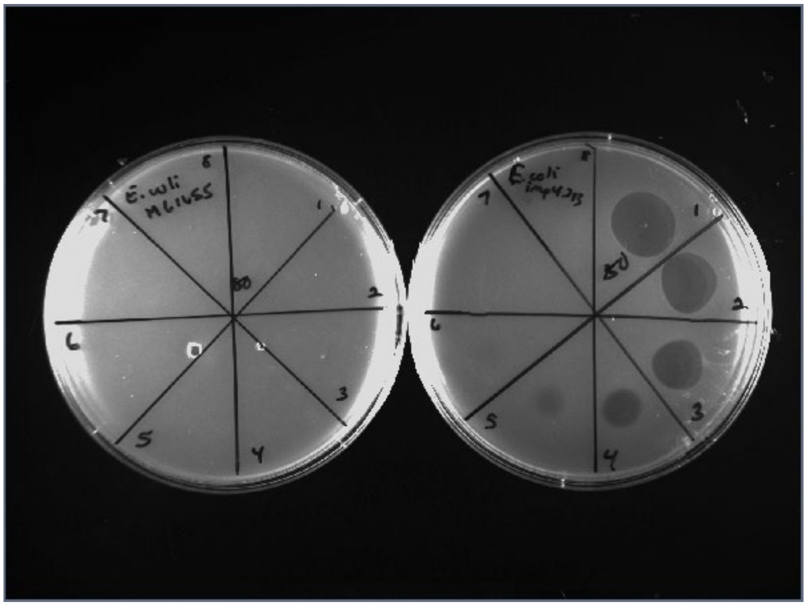

**Figure S13.** The antimicrobial activity of achromonodin-1 largely depends on its ability to traverse the outer membrane. Left plate: spot-on-lawn assay of achromonodin-1 against *E. coli* MG1655 in M63 agar. No activity is observed. Right plate: spot-on lawn assay of achromonodin-1 against *E. coli imp4213* in M63 agar. This strain has a compromised outer membrane, allowing for compounds to enter the cell that normally would be blocked by this permeability barrier. Achromonodin-1 has activity against this strain down to 2.5 μM. The highest concentration tested was 80 μM, and two-fold serial dilutions were spotted clockwise. The peptide was dissolved in pure water. Spot 8 is a pure water control.

**Table S1.** Chemical shift assignments for achromonodin-1 NMR in methanol. These assignments were used as input into CYANA 2.1.

| Residue | Hydrogen Atom | δ (ppm) |
|---|---|---|
| Glycine 1 | H | 7.491 |
| | A1 | 3.633 |
| | A2 | 4.445 |
| Glycine 2 | H | 9.262 |
| | A1 | 3.722 |
| | A2 | 4.446 |
| Glycine 3 | H | 8.250 |
| | A1 | 3.619 |
| | A2 | 4.332 |
| Glycine 4 | H | 7.718 |
| | A1 | 3.578 |
| | A2 | 4.487 |
| Proline 5 | A | 4.591 |
| | B2 | 2.239 |
| | B3 | 2.039 |
| | G2 | 2.132 |
| | G3 | 1.981 |
| | D2 | 3.406 |
| | D3 | |
| Threonine 6 | H | 7.716 |
| | A | 4.644 |
| | B | 3.301 |
| | QG2 | 1.228 |
| Proline 7 | A | 4.506 |
| | B2 | 2.154 |
| | B3 | 1.560 |
| | G2 | 2.063 |
| | G3 | 1.825 |
| | D2 | 3.573 |
| | D3 | 4.039 |
| Glutamic Acid 8 | H | 8.594 |
| | A | 4.100 |
| | QB | 1.626 |
| | QG | 1.880 |
| Tyrosine 9 | H | 7.724 |
| | A | 4.468 |
| | QB | 2.915 |
| | QD | 6.981 |
| | QE | 6.682 |
| Phenylalanine 10 | H | 7.867 |
| | A | 4.469 |
| | B2 | 3.066 |
| | B3 | 3.127 |
| | QD | 7.210 |
| | QE | |
| | HZ | |
| Leucine 11 | H | 7.826 |
| | A | 4.392 |
| | B2 | 1.549 |
| | B3 | 1.596 |

| | | |
|---|---|---|
| | G<br>QD1<br>QD2 | 1.501<br>0.767<br>0.800 |
| Methionine 12 | H<br>A<br>B2<br>B3<br>G2<br>G3<br>E | 7.900<br>4.698<br>1.947<br>2.046<br>2.486<br>2.553<br>2.026 |
| Proline 13 | A<br>QB<br>HG2<br>HG3<br>HD2<br>HD3 | 4.404<br>2.096<br>1.907<br>1.984<br>3.636<br>3.709 |
| Isoleucine 14 | H<br>A<br>B<br>QG1<br>QD1<br>QG2 | 7.687<br>4.192<br>1.883<br>1.495<br>0.913<br>1.169 |
| Aspartic Acid 15 | H<br>A<br>B2<br>B3 | 8.055<br>4.947<br>2.899<br>2.684 |
| Proline 16 | A<br>QB<br>HG2<br>HG3<br>HD2<br>HD3 | 4.353<br>2.280<br>1.881<br>2.000<br>3.836<br>3.949 |
| Alanine 17 | H<br>A<br>QB | 8.109<br>4.186 |
| Tryptophan 18 | H<br>A<br>B2<br>B3<br>D1<br>E1<br>E3<br>Z2<br>Z3<br>H2 | 7.733<br>4.402<br>3.310<br>3.402<br>7.232<br>10.390<br>7.360<br>7.109<br>7.024<br>7.497 |
| Leucine 19 | H<br>A<br>QB<br>G<br>QD1<br>QD2 | 7.621<br>4.076<br>1.660<br>1.582<br>0.867<br>0.930 |
| Glutamine 20 | H<br>A<br>QB<br>G2 | 7.965<br>4.072<br>2.098<br>2.340 |

| | | |
|---|---|---|
| | G3 | 2.374 |
| | E21 | 7.528 |
| | E22 | 6.844 |
| Alanine 21 | H | 7.817 |
| | A | 4.187 |
| | QB | 1.393 |
| Asparagine 22 | H | 7.809 |
| | A | 4.684 |
| | B2 | 2.752 |
| | B3 | 2.483 |
| | D21 | 7.284 |
| | D22 | 6.590 |
| Leucine 23 | H | 7.783 |
| | A | 4.467 |
| | B2 | 1.770 |
| | B3 | 1.564 |
| | G | 1.705 |
| | QQD | 0.929 |
| Proline 24 | A | 4.425 |
| | B2 | 2.286 |
| | B3 | |
| | G2 | 2.042 |
| | G3 | |
| | D2 | 3.754 |
| | D3 | 3.640 |
| Asparagine 25 | H | 8.316 |
| | A | 4.714 |
| | QB | 2.813 |
| | D21 | 7.658 |
| | D22 | 6.956 |
| Threonine 26 | H | 7.557 |
| | A | 4.449 |
| | B | 4.197 |
| | QG2 | 1.177 |
| Glycine 27 | H | 8.336 |
| | A1 | 4.042 |
| | A2 | 3.941 |
| Lysine 28 | H | 8.316 |
| | A | 5.206 |
| | B2 | 1.626 |
| | B3 | 1.716 |
| | QG | 1.174 |
| | D2 | 1.410 |
| | D3 | 1.508 |
| | E2 | 2.938 |
| | E3 | 2.980 |
| Tyrosine 29 | H | 9.235 |
| | A | 4.914 |
| | B2 | 2.998 |
| | B3 | 3.108 |
| | QD | 7.011 |
| | QE | 6.734 |
| Asparagine 30 | H | 8.087 |
| | A | 4.845 |
| | B2 | 2.741 |

| | | |
|---|---|---|
| | B3<br>D21<br>D22 | 2.859<br>7.576<br>6.804 |

**Table S2.** Explicit distance constraints used for achromonodin-1 model building in CYANA.

| Residue Number | Residue | Atom | Residue Number | Residue | Atom | Distance (Angstroms) |
|---|---|---|---|---|---|---|
| 1 | Gly | N | 8 | Glu | CD | 1.33 |
| 1 | Gly | N | 8 | Glu | OE1 | 2.26 |
| 1 | Gly | N | 8 | Glu | CG | 2.41 |
| 1 | Gly | H | 8 | Glu | CD | 2.06 |
| 1 | Gly | CA | 8 | Glu | CD | 2.42 |
| 1 | Gly | H | 1 | Gly | CA | 2.09 |
| 1 | Gly | CA | 8 | Glu | OE1 | 2.78 |
| 1 | Gly | H | 8 | Glu | OE1 | 3.17 |

**Table S3.** NOEs observed between steric lock residues and ring residues.

| Position | Residue | Hydrogen | Position | Residue | Hydrogen |
|---|---|---|---|---|---|
| 28 | Lys | HE3 | 2 | Gly | H |
| 28 | Lys | H | 8 | Glu | H |
| 28 | Lys | HA | 8 | Glu | H |
| 28 | Lys | QG | 8 | Glu | H |
| 28 | Lys | H | 3 | Gly | HA2 |
| 28 | Lys | H | 1 | Gly | HA2 |
| 28 | Lys | HB3 | 4 | Gly | H |
| 28 | Lys | HB3 | 6 | Thr | H |
| 28 | Lys | HB2 | 4 | Gly | H |
| 28 | Lys | HD3 | 4 | Gly | H |
| 28 | Lys | HD2 | 4 | Gly | H |
| 28 | Lys | HD2 | 1 | Gly | H |
| 28 | Lys | HE3 | 1 | Gly | H |
| 28 | Lys | HE2 | 1 | Gly | H |
| 28 | Lys | HB3 | 1 | Gly | H |
| 28 | Lys | QG | 1 | Gly | H |
| 28 | Lys | HA | 7 | Pro | HA |
| 28 | Lys | HA | 8 | Glu | QB |
| 28 | Lys | HB3 | 5 | Pro | HA |
| 28 | Lys | HB3 | 7 | Pro | HA |
| 28 | Lys | HB2 | 7 | Pro | HA |
| 28 | Lys | HE3 | 1 | Gly | HA3 |
| 28 | Lys | HE3 | 2 | Gly | HA3 |
| 28 | Lys | HD3 | 3 | Gly | HA2 |
| 28 | Lys | HD2 | 3 | Gly | HA2 |
| 28 | Lys | HB3 | 8 | Glu | HA |
| 28 | Lys | HD2 | 3 | Gly | HA3 |
| 29 | Tyr | H | 7 | Pro | HA |
| 29 | Tyr | H | 6 | Thr | HB |
| 29 | Tyr | H | 8 | Glu | H |
| 29 | Tyr | H | 6 | Thr | QG2 |

| | | | | | |
|---|---|---|---|---|---|
| 29 | Tyr | H | 8 | Glu | H |
| 29 | Tyr | QD | 8 | Glu | H |
| 29 | Tyr | HB3 | 1 | Gly | H |
| 29 | Tyr | QD | 8 | Glu | HA |
| 29 | Tyr | QD | 6 | Thr | HB |
| 29 | Tyr | QD | 6 | Thr | QG2 |
| 29 | Tyr | QD | 8 | Glu | QB |
| 29 | Tyr | QE | 8 | Glu | HA |
| 29 | Tyr | QE | 6 | Thr | HB |
| 29 | Tyr | QE | 6 | Thr | QG2 |
| 29 | Tyr | HB2 | 1 | Gly | HA3 |

**Table S4.** Percent identity of each RNAP subunit for tested *Achromobacter* strains to their homologs in *E. coli* RNAP. The β and β' subunits are where the lasso peptide microcin J25 is known to interact with *E. coli* RNAP, and these subunits are the most conserved from *Achromobacter* to *E. coli*. All *Achromobacter* RNAP subunits are >98% identical to their homologs in the other *Achromobacter* species.

| | *A. pulmonis* | *A. animicus* | *A. insolitus* |
|---|---|---|---|
| α subunit | 60.12 | 60.12 | 60.12 |
| β subunit | 65.80 | 65.80 | 65.57 |
| β' subunit | 65.84 | 65.48 | 65.62 |
| ω subunit | 44.78 | 44.78 | 44.78 |

**Table S5.** gBlocks used for assembly of achromonodin-1 *BCD* and achromonodin-2 *BCD* genes with preceding promoter for microcin J25 *BCD.* The corresponding gBlocks for each peptide were combined via overlap PCR before cloning into the achromonodin-1 or achromonodin-2 expression plasmid.

| Name | Sequence |
|---|---|
| gBlock1_achromonodin1 | GCGTTTTTTATTGGTGAGAATCCAAGCTAGCCATCAATTAAGAAAAAAATTTAGCTT<br>GTAGATAAATTCAGAAGTTTTATTATTCCAATTGAGTGTAAAGGCATAACTACAGGA<br>GGGAGTGTGCAAAATGCCGCAGCGTACACCGGGCAAATCCGTGCGCCTGGCGCA<br>CTTTCGTGATGATCTGATCGTCCTGAATCTGACCGAAAATCGTTTTAGCCTTATTCC<br>GAACATTCCGGTGGAGCGTATTAACTCATTTCTGGCGTCAGCGGGCAGCACCGAT<br>GCGACGTTGTCTGCGGCGCTTAGCCAGCATGGCATTACCAATGATAAACTGCCGC<br>TGAGCGCGGTGGTGAGCAGCCTGGAAAATACCGGCCTGCTGGAGACCCGTTGGC<br>GTACCCCGTTTACCCCGAGCGTCCGTTACAGTTTCCCGCTGCTGTGCCGTAGCATT<br>TGGACCCTGACGATGGCAGGGCTGATGATTAAAAGCGGAGGGTATGCGCTGGTGG<br>TCAAAGCGCTGCGTCGTAATAGCCAAAACACCAATGTGACCGCGATTGGCAGCGT<br>GGGCACCGACGAAATTATGGCGCATATCAATAAAGTGTTTCTGTTTGATGTGAGCA<br>ATAATCGTTGCCTGACCTATAGCCTGACTCTGTGCCTGATTGCACGGCGTATGGGC<br>TTGCCGGCGAAACTGATTGTGGGCGTTCGTACCCGTCCATTTTTCAGCCATGCGTG<br>GGTAGAACTGGATGGCGTGATCATTAACGACGATCCGGAACTGCGTGAAAAACTG<br>AGCATAATATTGGAAACATGATGTTACTGATTTATAGCCATAACCAGAATCTGGGCA<br>TGGCACGGGTGAGCGCGGATCCTAGCATCTCGACCACTCGGGCAGCACTGACCAA<br>AGATAACTTGATTGCGCGTTACTCGGCGGATTGGTCGGAACTTCAGAATAACGATT<br>GTCGTATCTTGTATTACGGACCGACCTACATCAAGACACAGATTAACATTCAGCGT<br>GATGAATTTCTGGGTCCGCATTGGCTGGCCGCTGCACATCGGCTGGACGGCGCAT<br>ATCTGCTGATTCGTATATTTGATACCCACATTGAGATTTCTCGTAGCCTTTATCGTG<br>GCCTGGACATTTTCTATTTACGTCTGGATGGTCGTCTGATTTTTAGCACCGATTTTA<br>GCTATCTGTTTGAAATAAGCGGCGAGAATAGCAACAACTTAGATGACAAGTATTGC<br>CATGATTTCATTATCGACTCGCCGACTTTCTCGGGCTCGACAGCGTTCAGCGCGAT<br>TAAAAGCCTGCAGCTGGGTCAATCCATTCGTACCGATCTGGATAGCATAACCGAAC<br>TGAGCCCGCAGGTGCCACGTCACACAGGGGGGAATCCAATTGATATCCTGACTGA<br>GAAGCTGAGCCATCTGGCGCGTCAGTTTGATAGCGTGTATCTGTCCTTTAGC |
| gBlock2_achromonodin1 | TGATAGCGTGTATCTGTCCTTTAGCGGCGGTCTGGACAGCAGCGTGGTGTTTTATT<br>GCCTGAAACGTGCGGGCGTGAGCTTTACCCCGTTTCATGGCGTGAATACCACACC<br>ATACGCGGAAAATGAGCTGGCGTATGCGAAGGATATTGCCCAGCATTTTGGTATTG<br>AACTGCAAGTGCGCTTTGCCAAAAACAATATCGAGCAGGCATTTTTCGCGCCGAGC<br>ACCTTTACCAAGACTACCTATCCGACTCCGTTTGCGGTGGATATTTTCACTCATCCG<br>GCGAGCGAAGCGAGCCAGAGCCAACGCTCCGAAAGCGAATATACCCGTCGTCTTT<br>ACCTGACCGGCCAGGGCGGTGATCATGTGTTTCTGCAAAATCCGGGCTGGAATGT<br>CGGCATTGATGATTTGCTGAAATTCCGTATTGGAAAAAGCATTTCACAAATTAGCAA<br>TTACTGCCAGCTGAAAGGCGAAAATCTTTATGCGCTGTTGTATCGGACCATTAAGTT<br>GGCGCTGCATAAGAACTCGTGCAAAAAGATCCTGAGTCACACTCCGCCGTGGATTT<br>CCCCTCCTGAACGTAGTGATCCGTCCGACCTGCATTACTTACTGAAAAATTCGGAC<br>CCCCGTAGCGCGAAGTTTCGTCATATTTCCACCATTCTTGCGGGCCTGAGCGGATT<br>AGCGTGGGACGATATAGCGGAAAGCAGCACAGTGCATCCGATGCTGCTGCCGGAT<br>ATGATTGGCACCGTGCTGGATCGTCCGGTGGGAGAACTGTATACCGCGAAATACG<br>ACCGTATTCTGCTGCGCCAGGCGGCTTATCAGGCTGCGGGGCATAACTTTTCTTG<br>GCGTCAAACCAAGCGTTCCTCAGCGACCGACTATTTTACCTATTTTCGTAATTATCT<br>TACCGAAATTACCCAGTGGCTGGAGAGTGGCATCCTGGTTAATAACCTGCAAATCG<br>ATCTGAGCAAGCTGCGGCAGAGCCTGGAATTTAACGGCTATTATCATCTGAATAAT<br>GACTTCCCTGCGCTGATAAACCTGATTCAGTTAGAAGCGTACATACAATCCACACT<br>GCACTTTTCATCGAGTCATCGGAATAGTTATGCTAAATAGCTTCCTGCCGCAAGTG<br>CTGCTTAAGGGCGGTCTGCGTGGCAAATCGGCGCAGTATCTGTTCGGCGCGACCG<br>CCGTGACACTGATTAGCAGCATTCTGAGCGCGGTTGCGCCACTTCTGTTAGCGCG<br>TATAACGAATGCCTTCAGCCATGCGGAAAATACCGCTGTGCTGTATGATAGTATTGT<br>GGTCCTGGGCGCGATATACATTTGCTTTGTGGCGTTTACCAAACTGCTGAACACCG<br>CGAGTTTGTACCTGCAGAGCCTGTTACGTATTCAGTTTATTGAAACCGTGAGCGAA<br>CGTTATTTCACCTATCTGTGCGAACGGGATGCGCAGTTCTTTGTGACCCACAGCGT<br>GGGTTCCCTGGCGCAGCAACTTAATCAGGCGACCAATGATC |
| gBlock3_achromonodin1 | GCAACTTAATCAGGCGACCAATGATCTGTATACCATTGTTCGGAATGTGGCCTTCA<br>ACATAGGCGCTCCCCTGATACAGCTGACCATTGCGATTTTGGTGATTACCAGCCAG<br>TTAGGCGGCGTGGTTGGCCTGATCTTCTTTCTGTATGCGGTATCCTTTTTTGTGAAT<br>AATCGCGTGTTTCTTAAGGAACTGAGCCCTCAGCATACACTGGTCATGGAAGCTGG<br>CCGTAAATCATATTCCACCCTGGTGGACTCCGTGATCAATATTACCGCAGTGAAAC<br>AATATAACGCCTTTAGCATTTTCTTTAAACGTTACAAGAATACATTGACCAGCGATC<br>GGAACATTCAGAATCGTTATTGGCGTCTGACCTTGACTATGCTTACCATTAATTCCC<br>TGTTGTTTGTGGTAATGTTTGCGCTGAGCGTGTATTGGATTCTGCATCATCCTAAAA |

| | |
|---|---|
| | ATGGAATCCTTACAGCGGGCGACTTCGTCCTGATTGCGACCTATGTGGTGCTGTTG<br>TCAGGCCCGATTGAATTGCTGGGTTCTACCTTTGGCGAAGTGGTACAGAGCTGGC<br>ATAGCGTCCGTCGTTTCCTGAGTGATACCTTTCAGAGCAAATTGTCCGCACCCGAG<br>ATTCAGCCGCTGCCGCGTAGCGGTGATGCGATTGTGCTGGAAGACGTCCTGTTTA<br>GCTATCGTAGCAGCCAGACCGTGCTGGGCCCTATTAGCTTCAAGATTAAAGCCGG<br>CGAAAAAGTGACCATCACGGGGAAAAGCGGAGCGGGTAAATCTACTGTGGCTCGT<br>CTGCTGACCGGAGAATATCCGCCGAGCCAGGGCAAAATTCGTATACATCAGCGGA<br>ATATCGAAGAAATTAGCCGTTACAGCTTGAATGAACTGATTGGTGTGATATCGCAG<br>GATATTTGCATATTCAATGATACCCTTCGTTTTAACATGCAGATAGCGGACGCGAAT<br>GCGACGGATGAGCAGATTGTGGAAAGCCTGACCCTGTCAGGGATGCCGATCCAAC<br>CATATGTCCTGGATCTGGATACACAATTGGGCGACCGTGGTGTGTCACTGAGCGG<br>TGGCCAGCGTCAGCGTATAGCACTGGCGCGCATTTTTCTGCGTAATCCGAGCATTG<br>TCATTATTGATGAGGGCACCTCCTCTCTGGATGTGAGCACCGAAAAGAGCATTAGC<br>CAAAACCTTTACAGCCGTTTTAGCGATTGCACTATTATCTCTATAAGCCATCGGCTG<br>TCGGCAATGCGGTTTTCTGAAAAGATTATCGTGCTGAAAAACGGAACCATTGAAGA<br>TAGTGGGACCATGCACGAATTGACCCAGCGTAATGAGTACATCAAAAACATGGTGA<br>AAATATCGTCTGAACAGAACCCGGGCTGACCATGGGCAAATATTATACGCAAGGCG<br>AC |
| gBlock1_achromonodin2 | GCGTTTTTTATTGGTGAGAATCCAAGCTAGCCATCAATTAAGAAAAAAATTTAGCTT<br>GTAGATAAATTCAGAAGTTTTATTATTCCAATTGAGTGTAAAGGCATAACTACAGGA<br>GGGAGTGTGCAAAATGCCGCAGGGCAATAGCGCGCGTGGCATTCAGGAACTGGT<br>GCTGGGCCATTTTCGTGAAGATCTGGTGGCGCTGGATCTGCAGGAAAATCGTTTTA<br>GCATTATTCCGAATCTGCCGCTTGCGCGGATTAATGAGTTTCTGGCGAATCGTGGT<br>CAGCATGATTCAGCCCTTGCGGCAGCACTGCAAGAACATGGCCTGCTGGAAAAAA<br>CCATTCATATTCGTGGCGAACTGGGCAGCCATATCAATGAAGGCCTGTTTGAAACC<br>CGTTGGGATACCCCGTTTATAGAATTTAGCCGCGCTGAACCGCTGTTGCTGTGGAA<br>AGCGGCGCATACCCTGTGGCGTGTGGGCTTAACCCTGAAAAGCAAAGGCTATGCG<br>GGCGTGATTAAACTGCTGCAGAATACCCAGCCTGGCCCACAGACCGCTCATACCG<br>CGAGCTATTTGCAGCGTCTGATGGCGCATCTGAATCGGGTGTTTATACTGGACTTT<br>TCCAACAATAAATGCCTGGCGTATTCTCTGGCGTTGTGCGTGCTGGCGCGTCGTGT<br>TGGCCTGGATGCGCGCCTGATAATAGGCGTCCGTACCCGTCCATTTAGCTCCCAT<br>GCGTGGGTGGAATTTAACGGCGCAGTGGTGAATGATGATCCGGAACTGCGTCAGA<br>AGCTGGCGGTAATTTTGGAACTATGATGTTTATTGCTCATTCGTCTGCGTTGCGTCG<br>TGATAATTGCACCCCGCGTAATGAACCTCTGACCTCTCATGGGAAACGGGCGATAC<br>ACGAGAACATTAATCTGGAATACCCGGATAACTGGAAGTTTATAGAAGATAGCGCG<br>ATTTCCATACTGCTTCATGCGCCCAAATATGTGGAAGAACAGCTGATAAATGCGCA<br>GAGCAAATTACGTACCGCGGAATGGCTGAGCTATCTGCACACTGTTGATGGCCCG<br>TATCTGGTGGTTCGTATTAGCTCGGGCAGCATTGAGATTAGCCGTAGCCTGTACCG<br>TGGACTGGACGTCTTTTATACCCATATTGCAGGATCCCTTGTGTTCAGCACTAGCC<br>CGTCCGTGTTACTTCGTTTAAAACGGGAAAAGAGCCATAGTTTAGACCTGGATTATT<br>GCCGTCGTTTCATTTTAGATCGGCCGGTGTTCTCAGGCTCCACGGCGATTGCGGG<br>CATCCGCGCGATACCGATTGGGCAAAGCATACGTAGCAATCTGGCAAGCATCTCTA<br>TAATTGGGTTACAGAAGCCACTGCCGCAACAGAAAGGCCCGATTGAAATACTGACT<br>AAAAAACTGAGCACTCTGGCGGCATCCTTTAGCAGCATCAGCTTATCATTTAGCGG<br>TGGCCTGGACTCCAGCGCGTTATTAC |
| gBlock2_achromonodin2 | CTGGACTCCAGCGCGTTATTACATTGCCTGGCGACAGCGAGCATTACCTTTGATGC<br>TATTCACGGCGTGAGCCCACTTCCGTATGCAGACACCGAACGCCGGGAAGCGGAG<br>CAGGTAGCCGCAGCGTACGGCAAACATCTGAAGCATATAAAGATGATCGAAAGCG<br>ATGCCCTGACCTTTACCCCGTCTTGCCTGAAGGAGCCGGCGTATCGTTCGCCGTT<br>CGACGTGAACATTTTTCTGACACCAGGCGCTAATCCTCAGCAGCAACAGGCGGTA<br>GCGGATTTCTCGGCGAATGAACTGATGATTACCGGTCATGGCGGGGATCATGTGTT<br>TCTGCAAAATCCGAATTGGAATGTGGGCTTCGACCGCATTCACAAACTGGAGGTTC<br>TGGGCTTTTTTCACGATGTGAAGCGTTTCTGTACCCTGAAAAAGGCGAACTTTTATC<br>GTGGAGTGCATGAAAACCTGCGCCTGCTTCTGAATCCACGGGGTTCTAATGGCGG<br>ATTGCATACCCCGAAATGGATGCAGGCGGGAGAAATTCTGCCTGCTGCCGACAAC<br>CATTATCTGCTGAATGAATTAGATCCACGTAGCGCCAAATTCGACCACGTATGGTCT<br>ATTCTGCTGGGCCTGAATACCATCCAGATGGAAAATAGCGTGGTGCGGACCACCAT<br>ACACCCGTTACTGTTGCCGGATGTCATTGGCACCGTGATGTATCGTCCGGTGCGT<br>GAACTGTTTAGCGCGCAGCATGATCGTTTTTACTTTCGTCGCGATCTGTATAAATCT<br>GCGGGCTATGACTTTGCGTGGCGTAAAAGCAAACGTAGCTCCTCCGCGTCTCTGTT<br>TAACTACTTTTCGGAAAACCATGCGTCCCTGTTCACCTGGCTGGAAGATGGCTTCA<br>TTGCAAAACAACTGACCCTGGATCTGGCCGAGTTAAAGAAGTCACTGAAGAAAAAC<br>GGTTCGACCTACTTGGATGATGACTTCCCTGCGCTGATTAACCTGATCCAGTTGGA<br>AGGGTTTGTGCAAAGCGTACTGCACCATAGTCAGTAAGGGCATACAAACATGAAAA<br>ATGGCAGCTTTCTGCTGCTGATACTGTCAGGCGGCCTGGGACATCGTTCGACCCTT<br>ATCTTTATGTCTGCACTGTTACTGACCCTGGCGTCGGCGGCACTGAGCAGCCTGG<br>CGCCTCTTTTGCTGGCGACTGCGACCGATATATTGGCTGGCAGCCAGAACATCGC<br>CGATACAATGTCCAGTGTCGCTTGGATGGCTGCGCTGTATATATGCAGCATTGGCT<br>TTACCAAAATCTTTAATACCATCAGCCTGTATCTGCAATCGATGCTGCGTTTGCAGC |

| | |
|---|---|
| | TTGTTGAGGCAATTTCTAAGAAATACTTTGCGCTTCTGTGTGAAAAAGACAGCCAGT<br>TCTACGTTCGGAACTCTCTGGGCGAACTTGCTCAACGTCTGAATCA |
| gBlock3_achromonodin2 | CGAACTTGCTCAACGTCTGAATCAGGCGTCAAACGATCTGTACACAGTCGTGCGTA<br>ATATTGCGTTTAACATCCTGGCACCGGCGGTCCAGCTTATTTTCGCAGTGTTTATTG<br>TGTCCAACATTCTGAGCGGCATTATTGGCATTGTTTTTTTGATCTATATTCTTCTGTT<br>CCTGATTAACAATCACGTGTTTTTGGACCGTCTTTCAAGCCATCGTACCCGTGTAAT<br>GGATGCGGGACGCCGTAGTTATGGCGTGCTGATTGATAGCGTGATGAACATTGTG<br>GCAGCGCGTCAGTACAACGGCTTTGACATTCTTATGAAACGTTATAAGACCACCTT<br>GGACGATGATCGGGCGGTGCAAAAAACCTACTGGCAGTTAACCATTACCATGTTAA<br>GCGTGAACAGCCTTCTTTTTGTCCTGATGTTCGGCTGGTGTATCTATTGGATGCTG<br>ACGGGGAGCAAGGACACTCAGACTACCCCAGGGAATTTTGTGCTGTTTGCGAGCT<br>ACATCTTGCTGTTGGCCGGACCCATTGAGATGCTGGGAAGCACCCTTAGCGAAGT<br>GCATCAGAGCTGGCACAGCGTTGCGGCGTTTGTGAAAGAACTGTTTCAGGGCAAA<br>TCCACCAATCAATCGCGTCCGAGCCTTGAAGCGGATGCTTATTCTGTGGAACTGGA<br>TCAGCTGGAATTTGATTATCCCGATGCAGCGCATTTTCATTTTGGGCCGATTTCCCT<br>GTGCTTTAAAAGCGGCGAAAAGATCGCCTTACTGGGCACCTCGGGAAGCGGCAAA<br>AGCACGTTGGCGAAACTTTTGACCGGCGACTATGCGCCTAGCCGTGGCGCTATTC<br>GGATCCTTGGCAATGACACAAGCGCGGTGTCACAAGCTAGTTTAAATGAACTTATT<br>GGAATTGTGAGCCAGGAAACCCACATATTTAGCGATACCGTCCGTTTCAACATGCA<br>GATTGCGGACTCGTCCGCGAGCGATTCTGAAATTCTGCGTGCGTTGGACCTGGCG<br>GGATTTGATGTCGCGAACAAGGACGGTCGTGAAAACATAAGCCTGGACGCTCAGT<br>TGGGAGAACGCGGCTTAAAGTTGAGCGGAGGCCAGCGGCAGCGTCTTGCGTTAG<br>CGCGCCTGTTTCTGCGCGAACCACGTATACTGATAATAGATGAAGGCACGTCCTCG<br>CTGGATGTGTTGGCAGAGCGTCGTATTACCGATAATATCACACGTCATTTTCGTAAT<br>TGCACCATCATTAGCATATCGCATCGCACGAGCGCACTTGCGTATAGCGAACGGGT<br>AATTGTTATTCATAATGGGAAATTACAAGCCGATGGCCCGAAGACCCAAATTGCGC<br>TGAGCAATGATTACCTGAAAAGCATGATGGCTATGTCTAACATAACCAAATGACCAT<br>GGGCAAATATTATACGCAAGGCGAC |

**Table S6.** Oligonucleotide sequences used to assemble *A* genes for both achromonodin-1 and achromonodin-2. These were assembled and cloned into the expression plasmids for achromonodin-1 or achromonodin-2.

| Name | Sequence |
|---|---|
| Achromonodin1_oligo1 | CACACAGAATTCATTAAAGAGGAGAAATTAACTATGCAGACCA |
| Achromonodin1_oligo2 | ATCGCTCTGAATCGGACGGGTGGTCTGCATAGTTAATTTCTCCTC |
| Achromonodin1_oligo3 | CCGTCCGATTCAGAGCGATAGCGAAATTCAGATTCTGAGCATTGC |
| Achromonodin1_oligo4 | GTCAGTTTGGTAGCCGGGGTGGCAATGCTCAGAATCTGAATTTCG |
| Achromonodin1_oligo5 | CCCCGGCTACCAAACTGACCCAGGGCGGTGGAGGGCCTACCCCGG |
| Achromonodin1_oligo6 | CGCCGGATCAATCGGCATCAGGAAATATTCCGGGGTAGGCCCTCC |
| Achromonodin1_oligo7 | TGCCGATTGATCCGGCGTGGCTGCAGGCGAATCTGCCGAATACCG |
| Achromonodin1_oligo8 | CTAATTAAGCTTTCAGTTGTATTTGCCGGTATTCGGCAGATTCGC |
| Achromonodin2_oligo1 | CACACAGAATTCATTAAAGAGGAGAAATTAACTATGCAGCG |
| Achromonodin2_oligo2 | GTTGCGGCTGCAGAATCGGACGCTGCATAGTTAATTTCTCCTCTT |
| Achromonodin2_oligo3 | CGATTCTGCAGCCGCAACCGCCGACCATTCAGGTGATTAAACTGG |
| Achromonodin2_oligo4 | TCAGTTTGCTCGCCGGAATTTCCAGTTTAATCACCTGAATGGTCG |
| Achromonodin2_oligo5 | ATTCCGGCGAGCAAACTGACCCAGGGCGGCAGCGGCGGTATTCCT |
| Achromonodin2_oligo6 | CCATAGGATCCGGCGCATAAAAGTATTCAGGAATACCGCCGCTGC |
| Achromonodin2_oligo7 | TTATGCGCCGGATCCTATGGGCTGGAAAAATCCGAATGTGACCA |
| Achromonodin2_oligo8 | CTAATTAAGCTTTTAGCCATATTTGGTCACATTCGGATTTTTCCA |

References for supplementary information


1. Hoover, D. M.; Lubkowski, J. DNAWorks: An Automated Method for Designing Oligonucleotides for PCR-Based Gene Synthesis. *Nucleic Acids Res.* **2002**, *30* (10), e43–e43. https://doi.org/10.1093/nar/30.10.e43.
2. Maksimov, M. O.; Pelczer, I.; Link, A. J. Precursor-Centric Genome-Mining Approach for Lasso Peptide Discovery. *Proc. Natl. Acad. Sci. U.S.A.* **2012**, *109* (38), 15223–15228. https://doi.org/10.1073/pnas.1208978109.
3. Madeira, F.; Pearce, M.; Tivey, A. R. N.; Basutkar, P.; Lee, J.; Edbali, O.; Madhusoodanan, N.; Kolesnikov, A.; Lopez, R. Search and Sequence Analysis Tools Services from EMBL-EBI in 2022. *Nucleic Acids Res.* **2022**, *50* (W1), W276–W279. https://doi.org/10.1093/nar/gkac240.
4. Güntert, P.; Buchner, L. Combined Automated NOE Assignment and Structure Calculation with CYANA. *J. Biomol. NMR* **2015**, *62* (4), 453–471. https://doi.org/10.1007/s10858-015-9924-9.
5. Hanwell, M. D.; Curtis, D. E.; Lonie, D. C.; Vandermeersch, T.; Zurek, E.; Hutchison, G. R. Avogadro: An Advanced Semantic Chemical Editor, Visualization, and Analysis Platform. *J. Cheminf.* **2012**, *4* (1), 17. https://doi.org/10.1186/1758-2946-4-17.
6. Cheung-Lee, W. L.; Parry, M. E.; Cartagena, A. J.; Darst, S. A.; Link, A. J. Discovery and Structure of the Antimicrobial Lasso Peptide Citrocin. *J. Biol. Chem.* **2019**, *294* (17), 6822–6830. https://doi.org/10.1074/jbc.ra118.006494.
7. Cheung-Lee, W. L.; Parry, M. E.; Zong, C.; Cartagena, A. J.; Darst, S. A.; Connell, N. D.; Russo, R.; Link, A. J. Discovery of Ubonodin, an Antimicrobial Lasso Peptide Active against Members of the *Burkholderia cepacia* Complex. *Chembiochem* **2020**, *21* (9), 1335–1340. https://doi.org/10.1002/cbic.201900707.
